# Flux-Balance Based Modeling of Biofilm Communities

**DOI:** 10.1101/441311

**Authors:** T. Zhang, A. Parker, R.P. Carlson, P.S. Stewart, I. Klapper

**Affiliations:** Department of Mathematical Sciences, Montana State University, Bozeman, MT; Center for Biofilm Engineering, Montana State University, Bozeman, MT; Department of Mathematics, Temple University, Philadelphia, PA.

## Abstract

Models of microbial community dynamics generally rely on a sub-scale model for microbial metabolisms. In systems such as distributed multispecies communities like biofilms, where it is not reasonable to simplify to a small number of limiting substrates, tracking the large number of active metabolites likely requires measurement or estimation of large numbers of kinetic and regulatory parameters. Alternatively, a largely kinetics-free methodology is proposed combining cellular level constrained, steady state metabolic flux analysis with macro scale microbial community models. The methodology easily allows coupling of macroscale information, including measurement data, with cell-scale metabolism. Illustrative examples are included.

## 1 Introduction

The study of microbial community dynamics has in large part employed tools of classical population ecology. Even with the increasingly sophisticated methodologies that have become available though, classical methods may not be well suited for exploiting the new depth of microbiological coverage because of their reliance on ad hoc constitutive law-based kinetics and regulation of those kinetics which, effectively, attempt to pre-compute relations between metabolism and environment. Further, classical methods rely on large numbers of reaction kinetics and regulation parameters that are unmeasured and possibly effectively unmeasurable (because they are themselves functions of environmental conditions), also making usage of detailed metabolic information problematic. The bioengineering community has, in response to these difficulties, introduced largely kinetics-free formulations at the cellular level, termed metabolic pathway (or metabolic constraint-based) analysis [4, 5, 35, 44] based on genomics and other omics (e.g., transcriptomics) derived data. Here we present a framework to incorporate kinetics-free methods into full scale microbial communities, biofilm models in particular, in part building on related methods previously applied to compartment models of eukaryotic cells [6, 7]. The key revision is to replace classical kinetics functions entirely by cell-level steady state metabolic pathway models – this is where omics meets community.

We focus here on biofilms, communities of microbes living and interacting at close quarters in self-secreted polymeric matrices [9]. Individual organisms take up substrates as locally available and utilize them to extract energy and build biomaterial, and also releasing waste in the form of metabolic byproducts. Processing is accomplished by, essentially, metabolic assembly lines termed pathways. Many internal and external metabolites can be involved [23, 53]. In many instances, limits on transport rates of chemical substrates into the biofilm and of byproducts out of the biofilm are important [48, 49]. On the other hand, substrate fluxes in and out of cells appear as sinks and sources at the large scale. Thus microscale and macroscales become closely coupled and hence a biofilm model provides a good test of a multiscale method.

Whether in a biofilm or in another type of community, individual microbes, and communities of microbes generally have many available metabolic pathways, and have extensive libraries of enzymes and regulatory mechanisms available to deploy in response to local environmental conditions. From a modeling point of view, a consequence is that kinetics parameters are functions of environmental conditions (by a variety of mechanisms, e.g., enzyme affinities and copy numbers). Thus determining rate parameters for metabolic models is difficult as rates are responsive to internal and external concentrations of, in principle, many metabolites [29]. In response, two main modeling strategies have been employed.

First, at the community level, kinetic models can be replaced by empirical constitutive laws relating, for example, external concentrations of a few limiting substrate to, for example, biomass production rates. This strategy amounts to precomputing metabolic function and can be effective when metabolism is relatively simple. However, spatially distributed microbial systems, like gut or soil communities, or biofilms, are often rather complicated with different metabolic pathways and networks active in different locations and times. Thus, anticipating metabolic activity with empirical constitutive response functions is likely to be difficult for models meant to address the increasingly detailed data that is becoming available.

A second strategy, steady state metabolic analysis, has been applied to cellular level models of metabolism. The key idea is to assume that kinetic dynamics reach a quasi-steady state. This assumption is justified in the case that environmental changes are slow in comparison to time scales for metabolic systems to reach steady state, i.e., kinetics time scales are relatively short in comparison to system times, as is often supposed to be the case. Mathematically, the quasi-steady assumption amounts to constraining internal reaction flux vectors to the null space of the stoichiometric matrix (though thermodynamics considerations such as restriction of some reactions as unidirectional as well as reaction flux bounds may further restrict the viable solution space to a cone within the null space.) That is, solution vectors satisfy flux balance - total fluxes into and out of all internal metabolites must exactly balance. This balance condition is entirely independent of the kinetic rate parameters; hence there is no necessity to measure or estimate their values. However, the choice of solution is generally underdetermined, even with the extra constraints – cells still have freedom to deploy available resources to different metabolic pathways. Thus some additional selection principle is required in order to distinguish a particular solution. This principle must rely on that information which is available, namely internal reaction fluxes (internal and external concentrations and other environmental information are not accessible to a cellular level model). Environmental conditions are represented by imposed maximum allowable influxes of metabolites. A typical optimization strategy is to choose from the admissible flux vectors the one that maximizes biomaterial production, though other optimization principles are also employed.

Steady state metabolic analysis thus has the important advantages that it does not require values for kinetic rates and, via optimization, can avoid need for details of regulatory mechanisms which, like kinetic parameters, are problematic to characterize. A cost, though, is the requirement for a selection principle that can be evaluated without access to important system level data. In fact, metabolic regulation should be able to respond to external environmental conditions [28] Likewise, such models are limited in their capability to generate system level information and so limited in ability to inform larger length and time scales of the community level which is often of the goal for microbial ecology and dynamics models. On the other hand, a community scale model with constitutive metabolic laws does, as a matter of course, produce much of that information, but its metabolic component is ultimately tied to kinetic rates. Thus combining the two scales is thus complementary. Hence the interest in combining metabolic pathway analysis with community scale models.

Some efforts have been made in this direction based on implicit or explicit iteration between a community level model which updates local external chemical concentrations based on physical processes, e.g. diffusion, and given fluxes of chemicals into cells, and a cellular level model which updates metabolic fluxes based on new influx bounds arising from updated chemical concentrations [10, 14, 21, 22, 24, 33, 36, 42, 47, 51]. This approach, though a convenient way to utilize already developed metabolic flux models, has some disadvantages. First, as generally practiced, optimization targets are largely inherited from the underlying cellular flux modeling so external information may not be utilized beyond setting flux bounds, though there are variants that look to combine metabolic analyses in a multispecies community [50, 56]. Even excepting this issue, these methods are generally built around a particular flux modeling approach. Second, this iteration procedure often matches cellular and system time scales, and so may be inconsistent with the foundational principle of metabolic flux analysis that metabolism is at quasi-steady state with respect to the environment. Further, from a practical point of view, in many cases the environmental time scales are of interest so that computing on the typically much shorter metabolic relaxation time scale is inefficient.

Here we propose to follow the standard multiscale practice of computing on the longer, system scale using a steady state cellular level assumption. Thus it is then natural to incorporate steady state metabolic models. Many choices for cellular level and system level modeling strategies can be employed, and, as a bonus, all available environmental information can be made available to the cellular model and vice-versa. To demonstrate, we model biofilm (a slowly growing microbial community) dynamics and employ external, environmental information to inform a cellular scale metabolism encoded by an elementary flux mode representation. Biofilms are, generally, relatively complex systems from both and ecological and mathematical viewpoint and hence serve as a useful vehicle to demonstrate methods [55].

## 2 Kinetics Free Modeling

### 2.1 Biofilm Model

We briefly describe a general biofilm community model [26] consisting of a domain Ω, a set of *N* material species with volume fractions *X*_*j*_(**x**,*t*), *j* = 1,…, *N*, and a set of *M* dissolved (exterior to microbial cells) metabolite (or substrate) or other chemical concentrations *C*_*k*_(x,*t*), *k* = 1,…, *M*. The material species, which may be actual biological species or other biological groupings or other particulate materials e.g. water or inert organic material, satisfy *X*_1_ + *X*_2_ + … + *X*_*N*_ = 1. The aim is to describe how, within the domain Ω, chemicals and materials are produced/consumed and are transported. Note that Ω itself may change in time as a consequence of material changes, e.g. production of new biomaterial.

The dissolved concentrations *C*_*k*_(x,*t*) satisfy transport equations, typically reaction-diffusion equations of the sort

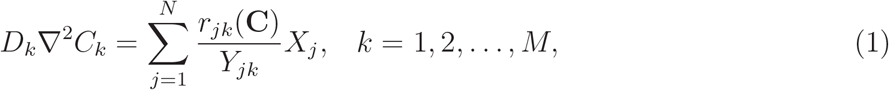

where *r*_*jk*_ is a consumption/production rate (units of time^−1^) and *Y*_*jk*_ is a yield coefficient (units of volume fraction per concentration) [26]. On larger scales, advective transport in a bulk flow may also be of importance and can be included [12, 18, 19, 37] but will not be considered here for simplicity. **C** is a vector of all dissolved concentrations exterior to cells. Note the assumption of quasi-equilibrium – for a relatively thin biofilm we suppose that the diffusion-reaction equilibration time is short in comparison to other system time scales such as biological growth. This assumption is again not critical but is common in biofilm models [37] and in some ways simplifies what follows.

The rate functions *r*_*jk*_ may typically be supposed to be empirically determined, for example based on a Monod product forms 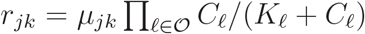 where, in this example reaction, chemical constituents are listed by index in 𝒪, half-saturations *K*_*ℓ*_ indicate uptake saturation of dissolved quantity *ℓ*, and *μ*_*jk*_ provides a maximum specific usage rate of dissolved quantity *k* by species *j*. Note that the complexity of system (1) can be expected to grow rapidly with increasing number of dissolved quantities in which case parameter estimation becomes problematic. In response, in many studies the vector **C** is restricted to a small number of “limiting” quantities, and neglected concentrations are assumed to be saturated. This simplification can be effective but may fail in spatially or temporally variable systems where it is not clear in advance which quantities should be considered the limiting ones at which locations and at which times. Even when a limiting approximation is appropriate, however, parameterization is still often uncertain.

Supplementing transport equations (1) for dissolved quantities, we also have dynamical equations for volume fractions of the form

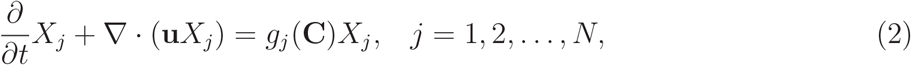

where *g*_*j*_ is a growth/decay rate rate and **u** is a material velocity. The rate function *g*_*j*_ may in fact be a function of the usage rates *r*_*jk*_, *k* = 1,2,…,*M* as, among these, some might relate to biomass production or maintenance. Note, also, we omit in (2) some effects that are sometimes included in biofilm models: conversion terms (e.g. conversion between active and inactive material), diffusive or other forms of mobility, etc. [15, 16, 27], as well as the possibility for different material types to move with different velocities [13, 25, 38].

Upon summing equations (2), we obtain the volume conservation constraint

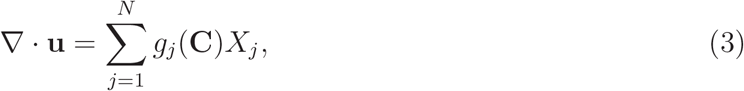

that is, the biofilm expands/contracts locally so as to accomodate local net material expansion/contraction. Later we will consider only variation in one spatial dimension (the coordinate *z* transverse to the biofilm) in which case, together with, say, a immovable wall condition 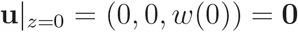 at a solid boundary, constraint (3) is sufficient to determine deformation the material everywhere. In more than one dimension, generally, system (2) is supplemented by a balance of forces equation in addition to constraint (3) in order to determine **u** [26]. Once **u** is determined then, at the free boundary *γ*(*t*),

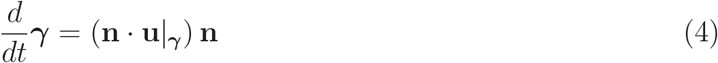

where **n** is an outward unit normal.

In what follows, we will limit to a single material species, i.e., *N* =1, and a short list of dissolved substrates (*M* = 3). In the case that *N* =1, the volume fraction *X* = *X*_1_ = 1 so that equation (2) becomes extraneous. Also, constraint (3) simplifies to ∇ · **u** = *g*_1_(**C**) = *g*(**C**) so that, for a one dimensional system, (4) reduces to

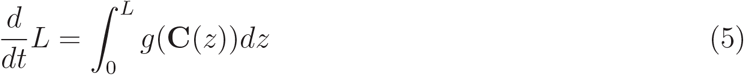

where *L*(*t*) is the biofilm thickness.

### 2.2 Metabolic Model

We, again briefly, introduce a general metabolic model consisting of the interior of a well-mixed cell, a set of m dissolved interior metabolite concentrations *c*_*k*_(*t*), *k* = 1,2,…, *m*, and a set of n reactions among the metabolites. Note that *m*, the number of internal concentrations, and *M*, the number of external concentrations, are not necessarily the same. A particular reaction can be represented by a chemical equation of the form

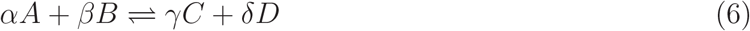

where *A, B, C, D* are metabolite labels (unitless) and *α, β, γ, δ* are stoichiometric coefficients (moles per reaction). The number of metabolites involved in any one reaction may of course vary from reaction to reaction.

The entire set of n reactions can be encoded in a mxn stoichiometric matrix *S* where the jth column of S corresponds to the jth reaction and the kth row corresponds to the kth metabolite. That is, if the kth metabolite is involved in the jth reaction, then *S*_*kj*_ is set to the corresponding stoichiometric coefficient (with minus sign if the metabolite appears on the left and plus sign if it appears on the right of the chemical equation). Otherwise, *S*_*kj*_ is set to 0. Given *S*, then

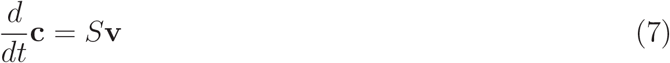

where **c** is the vector of interior metabolite concentrations (to be distinguished from the vector **C** of exterior concentrations) and **v** is the n-vector of fluxes (reactions per volume per time) through the n reactions. Generally, **v** = **v**(**c**) through a constitutive law which depends on kinetics, and kinetics parameters, of the metabolic reactions. However, using the steady state assumption of metabolic pathway analysis, equation (7) reduces to

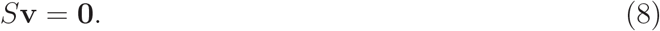

The idea is solve (8) for **v** rather than solve (7) plus constitutive law **v** = **v**(**c**) for **c**, as (8) is parameter free other than known stoichiometric coefficients in *S*, and knowledge of reaction fluxes is sufficient to characterize biomass production as well as influx and outflux of metabolites from cells. As a consequence though, internal metabolite concentrations **c**, even at steady state, are unknown.

However, there is a caveat: often there are more reactions than metabolites in which case system (8) is underdetermined. Hence some criteria must be adopted by which to choose a designated flux vector v from the solution space of (8). Much attention has been given to developing such principles and there are a number of options available; we do not need to assume use of any one such methodology here. Note that the choice of null space vector is not entirely unconstrained even before biological function is considered, as thermodynamical considerations make some vectors in the null space of *S* very unlikely to occur in practice [39, 40]. However, these restrictions involve metabolite concentrations c which are unknown so are generally difficult to apply. However, in some cases thermodynamic constraints can be imposed anyway, one important example being reactions that proceed in only one direction except at unrealistically extreme values of the reaction component concentrations, i.e., reactions for which the flux can effectively assumed to be of one sign only (with the reaction conventionally set so that its flux is nonnegative). Thus equation (8) is often supplemented by a number of non-negativity constraints of the form *υ*_*ℓ*_ > 0, where *ℓ* ranges over the set of effectively single-direction reactions.

In preparation for connecting to a macroscale community model, we replace (8) by a set of such equations

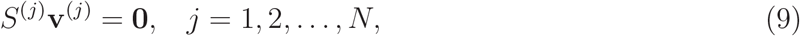

one for each of the N material species. That is, each active material species will have its own set of internal reactions and metabolites, and so its own stoichiometric matrix *S*^(*j*)^ and flux vector **v**^(*j*)^ (with *S* = 0 for inactive material species). Conversely, one can think of the stoichiometric matrix *S*^(*j*)^ as defining species *j*. Note that *S*^(*j*)^ is independent of **x**, whereas the distinguished solution **v**^(*j*)^ = **v**^(*j*)^(x) generally. Given, then, flux vectors **v**^(*j*)^, the growth rates *g*_*j*_ in the biomaterial equations (2) can also be replaced by parameter-free functions of biomass-producing (or removing) reaction fluxes, i.e.,

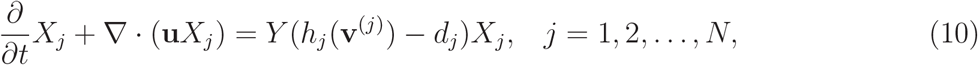

where **v**^(*j*)^ is the vector of fluxes involved with species j, *h*_*j*_(**v**^(*j*)^(**x**)) is the net specific biomass production rate for species j, *d*_*j*_ is a decay (or maintenance) coefficient, and *Y* converts biomass to volume.

### 2.3 Exchange Fluxes

Internal and external (to cells) chemical concentrations are connected through exchanges of the form

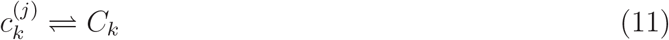

between internal concentrations 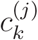 (kth substrate internal to species j) and external concentrations *C*_*k*_. When convenient, we can regard (11) to be a reaction, as in (6), in a general sense. Exchange reactions serve as the couplings between the cellular metabolic model and microbial community model. That is, if reaction (11) proceeds with flux *e*_*jk*_(**x**) (units of concentration per time per volume fraction) between interior and exterior of material species *j* at macroscale location **x**, then system (1) can be replaced by

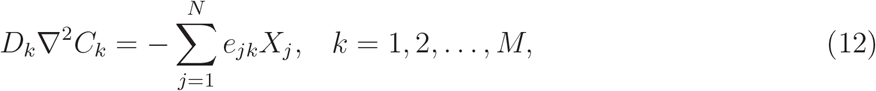

eliminating the need for the empirical rate functions *r*_*jk*_. Note that *e*_*jk*_(x) measures the specific flux of metabolite *k* produced by material species *j* into the exterior domain. Conversely, exchange reactions appear in the stoichiometric matrix *S* as mono-component reactions

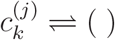

where the notation ( ) denotes exit from the internal cellular metabolic system. Each exchange reaction corresponds to a column of *S* containing all 0s except for a single 1 entry. The fluxes *e*_*jk*_ are then the components of solutions **v**^(*j*)^ to (9) corresponding to those same columns.

Fixing a flux for exchange (11) sets a constraint for the metabolic model: internal fluxes in and out of species *j* must balance this exchange flux. In practice, metabolic modelers do not necessarily fix exchange fluxes but rather might set upper bounds on them. We will do the same here. Choosing the value for an upper bound on flux into a cell is a physical problem depending on material transport properties of the particular species and also, possibly, biologically regulated transport mechanisms. Here we suppose that flux out of cells is unbounded. Thus, flux *e*_*jk*_ for exchange (11) would be required to satisfy inequalities of the form

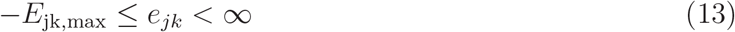

Then, in practical terms, all of the rate functions *r*_*jk*_ and their attendant parameters are replaced by a set of maximum realizable influxes *E*_*jk*,max_, commonly determined as functions of exterior concentrations **C**.

### 2.4 Optimization

Equations (9) for the metabolic models together with equations (10) and (12) for the macroscale biofilm model make up the foundations of a kinetics free system. As already noted, though, system (9) is underdetermined and requires a selection principle for choosing a particular flux vector out of the full solution space. Metabolic pathway flux analysis will typically make this choice based on a cellular level optimization principle such a maximum biomass production rate or maximum energy usage rate, though other methodologies are also possible [43, 45]. Such cellular level optimization principles can be used here as well, though we are free to also incorporate in part or in full external, environmental information.

As an example of an environmentally-based principle, and one that we will use below, we could choose flux vectors in order to minimize external resource availability in some prescribed manner, with motivation that an ecologically stable community would find itself in an environment which is sparse enough that invasion by outside microbes cannot succeed. As a particular measure of resource availability, we will choose the enthalpy of combustion (heat released when chemical constituents are fully combusted with oxygen) in the exterior environment; other measures are certainly possible. Combustion enthalpy provides a rough estimate of available catabolic potential and, for the simple metabolic models considered here, a proxy for growth potential.

Two additional notes: first, with multispecies and spatially distributed communities comes extra freedom. Different species in one location as well as species (different or not) in different locations will have different choices of flux vectors and perhaps different optimization goals. As one example, one could ask that all flux vectors are chosen in concert so as to minimize the system-wide objective using L_1_, or System, optimization. Alternatively, one could ask that independent bio-groups choose choose their own flux vectors so as to optimize their own, local objective, independently of strategies of other bio-groups. This latter strategy is, essentially, Nash optimization [34]: the system-wide flux vector profile is such that it is nowhere the case that a local perturbation would locally (only) improve optimality. Other minimizations methodologies, beyond System and Nash, are also possible, and in general different optimization schemes will in general lead to different results. An example of such is provided later.

Second, it is also possible to employ various sorts of optimization objectives. As one alternative, rather than (or in addition to) physically motivated objectives, fitting of empirical data might be used. Specifically, we can ask for flux vectors that result in system solutions that, in addition to or alternatively to resource optimization, best match a given criterion such as measured data. For instance, we can suppose that additional boundary concentration or flux measurements are available through observation, and ask that solutions of (9) are selected so as to produce model boundary concentrations or fluxes that best match these data, again see computations presented later for an example.

## 3 Example System

### 3.1 Metabolic Model

To illustrate, we consider a system consisting of a single species biofilm grown on top of an agar base (providing a nutrient source, supposed here to be glucose) with exposure from above to air or bulk fluid (providing an oxygen source). The microbe is provided with a simplified metabolism, see Figure 1, in which glucose (Glu) converts to biomass either through full respiration in combination with oxygen (O_2_) or, without oxygen, through fermentation. In the latter case, a fermentative product, supposed here to be lactate (Lac), is produced as a byproduct. Lactate itself can also be fully respired with oxygen to produce biomass. Note that this metabolic model is intended as a coarse-graining of a true metabolic map which would include, generally, many more internal metabolites and internal reactions.

**Figure 1:**
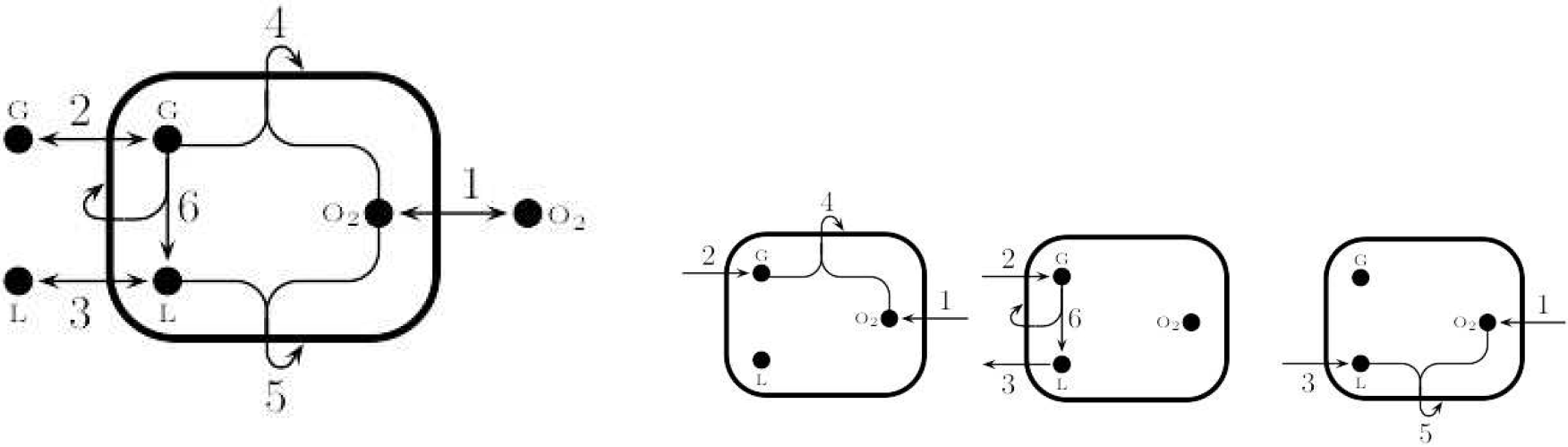
(Left) Metabolic model consisting of three metabolites: oxygen O_2_, glucose G, and lactate L, and six reactions: exchange reactions 1-3 connecting interior and exterior pools of oxygen, glucose, and lactate respectively, as well as interior paths for glucose respiration (reaction 4), lactate respiration (reaction 5), and glucose fermentation (reaction 6). Reactions 1-3 are reversible and reactions 4-6 are non-reversible. Hooked arrows indicate biomass production. Stoichiometric coefficient are given in (15). (Right) Elementary flux modes for the metabolic model on the left. From left to right: glucose respiration, glucose to lactate fermentation, and lactate respiration. See (16) for the corresponding flux vectors including stoichiometry.

For this system, in the notation introduced above, *N* = 1 and *M* = 3, and the exchange and internal chemical equations are

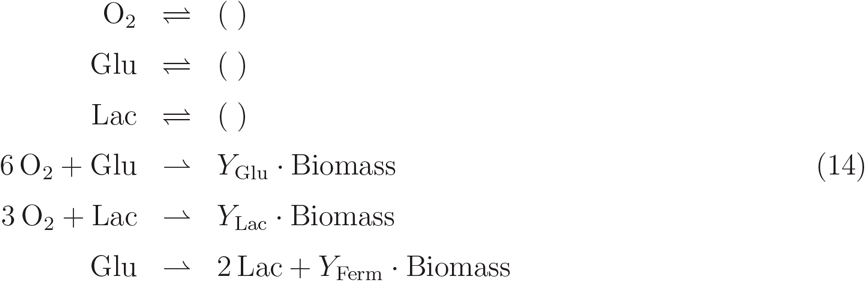

where the first set of three equations denote exchange between internal and external metabolites, and the second set of three equations denote internal metabolism. “Biomass” refers to generation of new biomass. The coefficients *Y*_Glu_ = 19, *Y*_Lac_ = 16, and *Y*_Ferm_ = 0.5 are yields (biomass per reaction), and help determine the rate at which given pathways generate biomass. Note that they are effectively arbitrary here, because time units are arbitrary so that rates of generation of new biomass are arbitrary. However, their relative values matter – here we have set glucose respiration to be a bit more efficient than lactate respiration, and both are significantly more efficient than glucose fermentation. Additional products CO_2_ and H_2_O are suppressed in (14), as is the nitrogen source, because they play no direct role here (for example, some carbon is lost from the system in the form of respired CO_2_ but this does not effect results below directly). Including them, and others, can allow the reactions to be elementally and electron balanced. We don’t attempt to represent any actual microorganism though; rather, the stoichiometry and yields are only intended to be illustrative. In fact, reactions (14) are heavily simplified in some ways-they implicitly connect ATP production with biomass yield and suppose another source of carbon for anabolism.

The stoichiometric matrix for reactions (14) and for the metabolic model in Figure 1 is

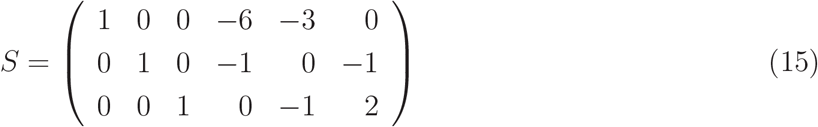

with the six columns encoding the stoichiometry of the six reactions above and as numbered in Fig. 1, and the three rows corresponding to the three interior metabolites oxygen, glucose, and lactate respectively. A row for “Biomass” is not included in *S*; rather, biomass-related stoichiometric coefficients enter into the external rate *h*, see equation (10) and below. A solution vector **v** of *S***v** = **0** has as its components fluxes through the six reactions. Some of the reactions are supposed unidirectional; we allow a solution vector v only if all of its unidirectional components are non-negative. For this example, thus, we require components 4-6 of v to be non-negative. We also follow convention and, for the reversible exchange fluxes 1-3, denote flux from interior to exterior as positive.

Graphically, a solution of *S***v** = **0** can be viewed as a pathway through the metabolic map. Examples can be seen in the elementary flux modes [8, 11, 46, 54], see Figure 1 right, for the example system. Elementary flux modes (EFMs) are a set of allowable pathways through the metabolic model (i.e., pathways that correspond to vectors in the null space of the stoichiometric matrix that satisfy non-negativity constraints) which are minimal in the sense that an EFM contains no sub-pathway which is also an EFM. Here, the set of EFMs (unique up to normalization) correspond to the three metabolic modes of glucose respiration, glucose fermentation with lactate byproduct, and lactate respiration. Denoting them by **v**_1_, **v**_2_, and **v**_3_, and normalizing by input carbon content, we can write them in vector form as

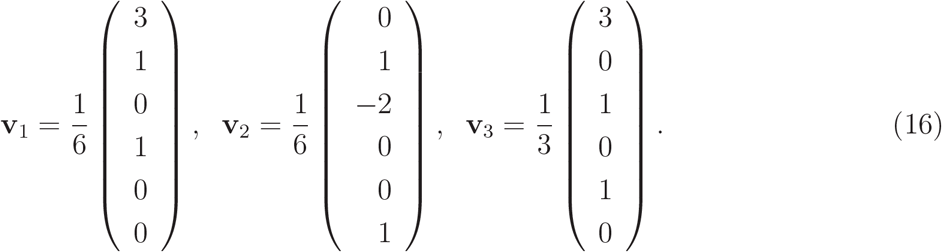

For a general metabolic map, any allowable pathway vector can be shown to be a (generally nonunique) non-negative linear combination of EFMs. For the simplified system here, it happens though that each allowable pathway vector does have a unique representation

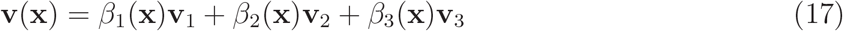

with flux amplitudes *β*_*j*_ ≥ 0. Note that the number of EFMs generally increases rapidly with the size of the metabolic model [23] making an EFM representation increasingly complicated in more detailed systems.

Underdetermination of (8) manifests here in the freedom in (17); any non-negative set of *β*_*j*_>’s results in an admissible solution **v**. Thus a scheme, including additional constraints such as (13), must be introduced to distinguish particular choices. This will be addressed below. Once chosen, however, the biomass production flux can be computed as *h*_*j*_(v^(*j*)^) = *h*(*β*), see (10). For the example, here

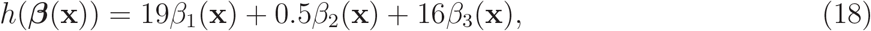

with yield coefficents indicating Cmoles of biomass produced per pathway reaction.

### 3.2 Exchange Fluxes and the Biofilm Model

In this system there are three exchange fluxes of the form (11) between internal and external oxygen (*c*_1_ ⇌ *C*_1_), internal and external glucose (*c*_2_ ⇌ *C*_2_), and between internal and external lactate (*c*_3_ ⇌ *C*_3_). These three fluxes are denoted, since this is a one species system (N=1), by *e*_1*k*_ = *e*_*k*_(**x**), *k* = 1, 2, 3. We bound inflow following (13), with *j* = 1, by

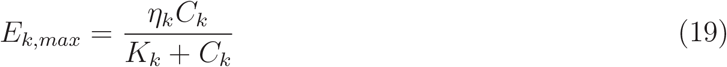

where *η*_*k*_ determines a maximum transport rate and *K*_*k*_ determines transport rate saturation.

In terms of the flux amplitudes *β*_*j*_, see (17), the exchange flux functions *e*_*jk*_ (**v**(**x**)) = *e*_*k*_ (**v**(**x**)), *k* =1, 2, 3, see (12), are given by the components *e*_*k*_(**v**) = −**v** · **i**_*k*_ of **v** corresponding to exchange reactions, where **i**_*k*_ is a 6-vector with a 1 in the kth place and 0s elsewhere. In this example,

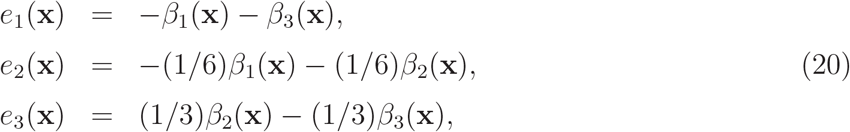

where the coefficients indicate pathway stoichiometry: moles of substrate *k* (here oxygen, glucose, and lactate) produced per pathway *ℓ* (here glucose respiration, glucose fermentation, and lactate respiration). In (20), *e*_1_ is flux of oxygen, *e*_2_ is flux of glucose, and *e*_3_ is flux of lactate (following the sign convention that *c* ⇀ *C* is considered positive).

With these specifications for exchange fluxes, the 1D biofilm model (1)-(5) reduces for this example to

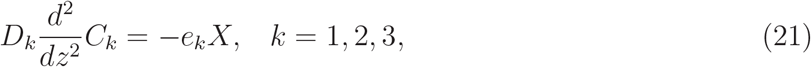

with *z* ∈ (0,*L*(*t*)) where, from (2), (5), and (10), the height/thickness of the biofilm changes according to

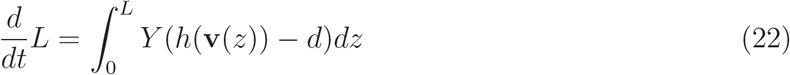

with *e*_*k*_ and *h* as described in (17), (18), and (20) above. Together with boundary conditions (given below), this system is complete up to determination of *β*_1_, *β*_2_, *β*_3_. A cartoon is shown in Figure 2: the righthand sides of (21) are represented, essentially, by horizontal arrows (exchange fluxes), and the lefthand sides are represented, essentially, by vertical fluxes (diffusive and other physical fluxes). In fact, physical fluxes could be considered as additional chemical reactions [17] and added in with the others, though such fluxes generally have well established constitutive relations with concentrations. The black rectangles in Fig. 2 are black boxes for purposes of the biofilm model; what happens inside is the domain of the metabolic model and is communicated to the biofilm model through exchange fluxes and biomass production. Biomass production results in pressure-driven expansion over long times, see (22), which causes the biofilm to gradually expand.

**Figure 2:**
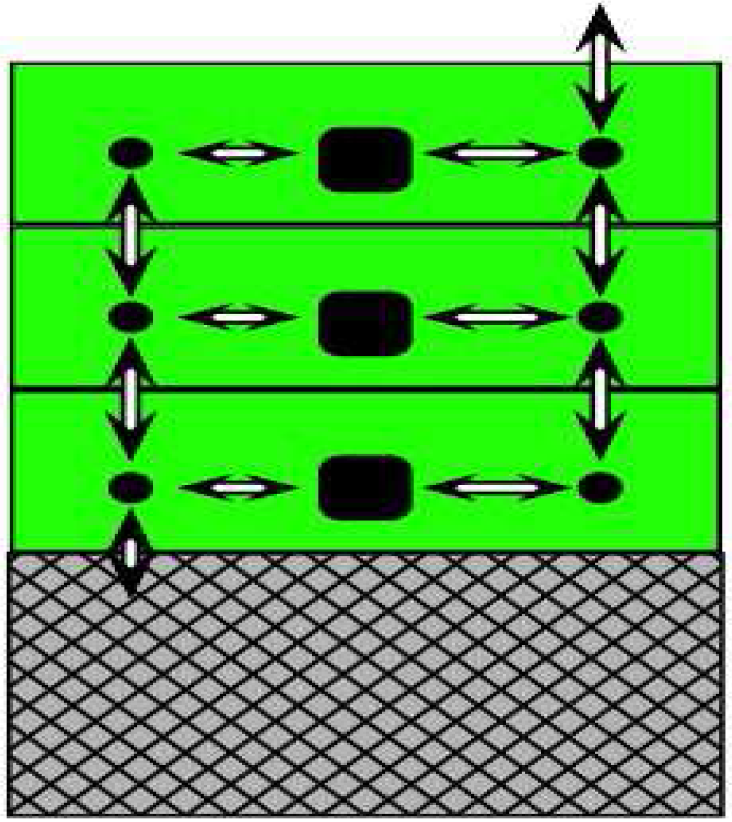
Macroscale biofilm (green solid) on solid substratum (gray hatched), open to air or bulk fluid above. Quantities vary in the vertical direction only, and each green layer corresponds to a different spatial location. Black boxes represent internal, microscale metabolic models (see Fig. 1) in different layers and black circles represent exterior chemical concentrations in different layers. Horizontal arrows are exchange fluxes; vertical arrows are diffusive fluxes. Chemical quantities may or may not be able penetrate into or out of the substratum (permeability) or into/out of the air (volatility), encoded in the boundary conditions.

To complete the model (biofilm plus metabolism), choice of a metabolic flux vector **v** satisfying *S***v** = **0**, with *S* given in (15), is required. This amounts to fixing values of the flux amplitudes *β*_1_, *β*_2_, *β*_3_, which are arbitrary up to the requirement that all constraints be satisfied. In this example, the three EFMs are all unidirectional, so we require *β*_*j*_(**x**) ≥ 0. We also assume that the exchange fluxes satisfy maximum influx constraints of the form (13). As *e*_*jk*_ = *e*_*k*_ here, with bounds *E*_*k,max*_ as formulated in (19), then constraints (13) become

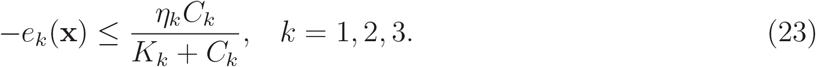

Note that the righthand sides of inequalities (23) depend on values of *β*(**x**) through the entire domain. Given inequality constraints (23), then, the optimization procedure is described next.

### 3.3 Optimization

Two issues need to be addressed: an optimization principle must be chosen, and a measurement procedure must be introduced. For the first, we apply physically-based optimization principles such as maximization of rate of biomass production or maximizing entropy production [20, 52]. Alternatively, a more directly biologically-based principle, like maximization of biomass production can be used. Typically in metabolic analyses, these principles are implemented at the interior cellular level. Here, however we can optimize at the cellular interior or exterior, or system-wide levels, or any combination. As an example, we determine *β* by minimizing external enthalpy of combustion, defined as the enthalpy released after complete combustion by oxygen. A possible motivation is ecological – in order for a community to be stable against invasion, it might reduce the available (i.e., exterior to cells) energy to as low a level as possible. However, we make this choice of objective for illustrative purposes without claiming it to be better or worse than other possibilities, and it is not the only possibility. External enthalpy of combustion density *H*(x) is given by

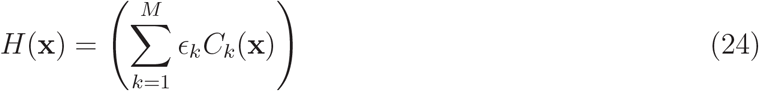

where *ϵ*_*k*_ is the standard enthalpy of combustion per mole for substrate *k*. In the example system, *M* = 3 with *ϵ*_1_ = 0, *ϵ*_2_ = 2805, and *ϵ*_3_ = 1344, in units of kJ/mol, for substrates oxygen, glucose and lactate respectively [31]. The vector *β*(**x**) is chosen to minimize a measurement of *H*(**x**).

In addition, we also present an entirely different type of optimization principle in which determination of *β*(**x**) is made so as to best match a given set of data. For example, given a profile of, say, *C*_1_(**x**), we can ask for a determination of *β*(**x**) such that the solution of the biofilm problem (at a fixed time *t* for simplicity) best matches that profile. The motivation comes from a sort of inverse problem: suppose that we know something of the genetic capability of a microbial population and, in addition, we have some measured environmental data, e.g., an oxygen profile. To what extent can we determine the metabolic activity of the system? At the same time, we may be able to predict unmeasured environmental data, e.g. the glucose and lactate profiles.

Whatever the optimization principle, some method of measurement of optimality is also required. Optimality of the argument of a spatially dependent function like that in (24) requires choice of some statistic. We illustrate with two different strategies. First, optimization is measured via Nash equilibration where the choice of *β*(**x**) is made at each location x without regard to its effects at other locations. That is, we consider 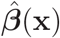 to be a Nash optimizer if, given any location x_0_, any allowable perturbation to 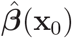 results in an increase in *H*(x_0_). Second, as an alternative strategy, we optimize over the entire biofilm by choosing *β*(**x**) to minimize 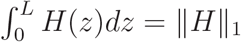. Here the entire microbial community works together to minimize the total, system-wide enthalpy of combustion. System optimization might be less biologically plausible than Nash equilibration (though perhaps there can be cooperation of some sort through signalling or other mechanisms [30, 32], and system wide optima might suggest consortial strategies). However, the main motivation is illustrative; again we make no claims as to what method is best and only intend to illustrate the possible.

## 4 Parameters and Methods

We present in Section 5 results of simulations based on the example system biofilm described in Section 3. Some details are described here, though.

### 4.1 Boundary conditions

First, boundary conditions for the system are chosen to be representative of what might be used to simulate a biofilm with tissue interface (*z* = 0) and air interface *z* = *L*, where *L* is the coordinate of the top boundary of the biofilm: for oxygen (*C*_1_), Dirichlet BC at the top *z* = *L* and at the bottom *z* = 0: for glucose (*C*_2_), Dirichlet BC at the bottom *z* = 0, no-flux BC at the top *z* = *L*: for lactate (*C*_3_), zero Dirichlet BC at the bottom *z* = 0, no-flux BC at the top *z* = *L*. In particular, at the bottom boundary *z* = 0,

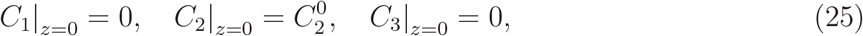

and, at the top boundary,

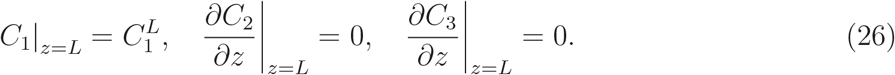

Effectively, glucose is supplied to the layer from below (*z* = 0) and oxygen from above (*z* = *L*). Lactate enters the system only internally via fermentation. It is assumed that glucose and lactate are unable to cross the upper boundary into the “air” region. At the bottom, the “tissue” interface, lactate and oxygen are removed from the system.

### 4.2 Model parameters

Lengths are scaled in units of 100 *μ*m, e.g. *z* = 1 means *z* = 100 *μ*m, times are scaled in units of hours, e.g., *t* =1 means *t* =1 hr, and concentrations are scaled by a reference concentration 4 × 10^−4^ mol · l^−1^, e.g., *C* =1 means *C* = 4 × 10^−4^ mol · l^−1^. See Table 1 for a list of parameter values used. In addition to parameter values already discussed elsewhere: diffusion coefficients are chosen based on measurements made in biofilm environments [48], conversion coefficient value *Y* is based an assumption for live cells of specific volume = 0.91 cm^3^·g^−1^. As the “microbe” we simulate is essentially artificially constructed for illustrative purposes, we don’t attempt to match parameters in constraint bounds (23) to the literature. Rather, values for these parameters are chosen to result in significant growth, for substrate concentrations present in simulations, on the designated system time scale of hours. Similarly, the decay rate *d* is not chosen based on literature reports but rather in order that the time scale for transition from thin to thick biofilm regime also occurs on that system time scale.

**Table 1:**
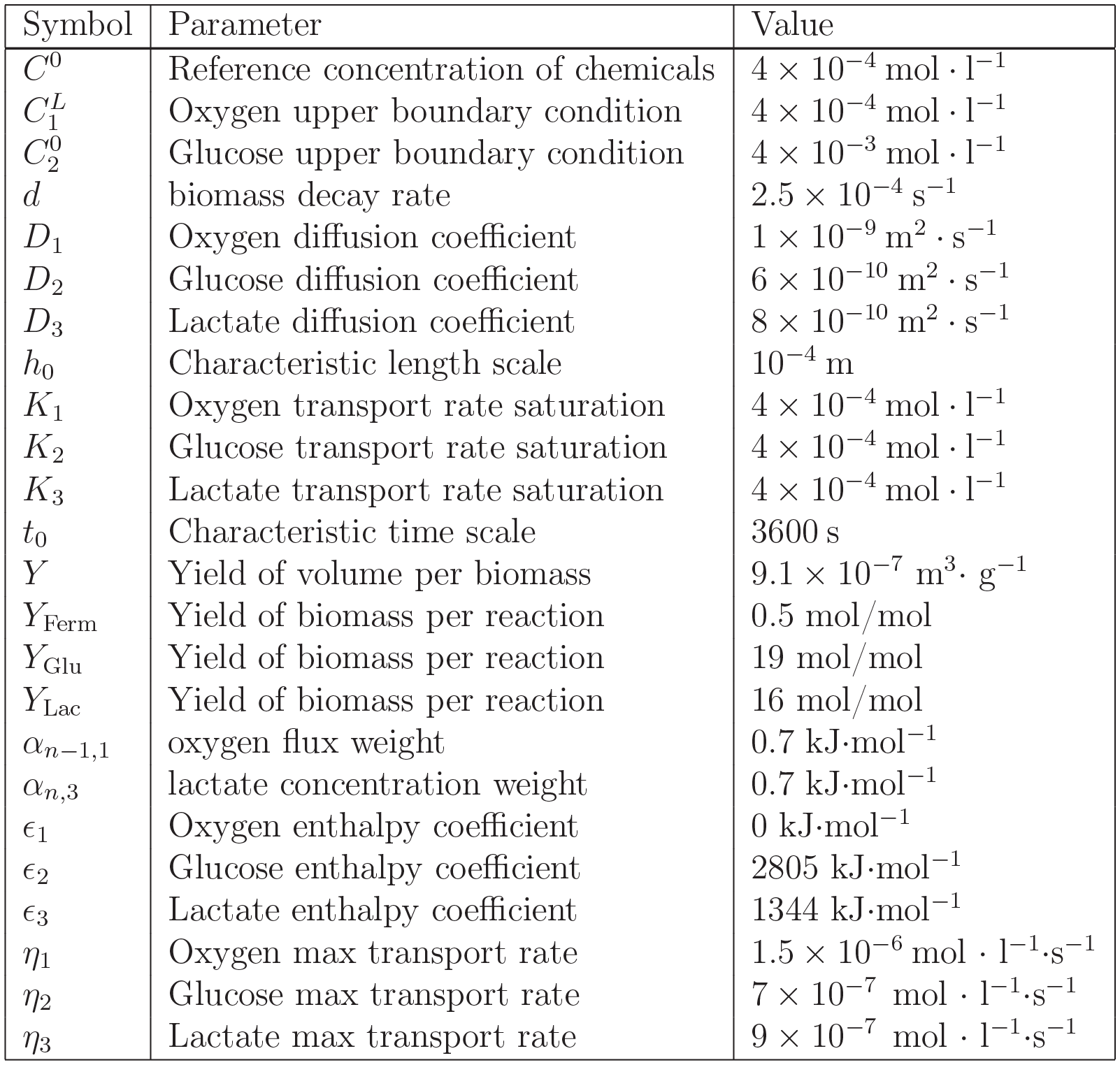
Parameter values used in computations. Other than *K*_*j*_s, *η*_*j*_s, and *d*, values are either free, given by stoichiometry, or measurable at the macroscale.

### 4.3 Computational details

Equations (21) are defined on a domain *z* ∈ [0,*L*] with a moving upper boundary at *z* = *L*(*t*). For convenience, a new spatial coordinate ζ = *z*/*L*(*t*) is introduced in order to obtain a fixed computational domain ζ ∈ [0,1], and equations (21) are written in terms of variable ζ, as in [55]. These equations are solved for *ζ* ∈ [0,1], and results are reported on the original spatial domain *z* ∈ [0,*L*]. Standard finite difference methods are employed with the spatial domain [0,1] discretized into n subintervals of size Δζ = 1/*n* (*n* = 20 in the computations shown below). Flux amplitudes and concentration values are tracked at the center of each subinterval, and denoted as follows: 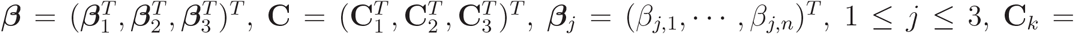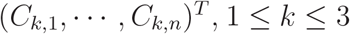. A given flux *β* determines the right hand side of equations (21), and the corresponding concentration profiles **C** are obtained by solving equations (21). We denote this mapping from *β* to **C** by a function **C** = *G*(*β*).

(1) For a fixed biofilm thickness *L*, the flux amplitudes *β* and the corresponding concentration profiles **C** within the biofilm are computed by solving the optimization problem discussed earlier, with details given below. (2) In the case of growing biofilm simulations, an initial biofilm thickness *L*_0_, a total simulation time *t*_*F*_, and a growth time interval Δ*t* are chosen. (Results shown below are for *L*_0_ = 0.1, *t*_*F*_ = 15, Δ*t* = 0.02.) Then, at each time *t*_*m*_ = *m*Δ*t* and biofilm thickness *L*_*m*_ (at that time), the flux *β*_*m*_ is computed by optimization as in the fixed thickness computations, the rate of change of the thickness 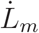 is computed by equation (22), and then the thickness is updated by 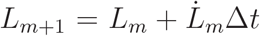. Time stepping continues until the total simulation time *t*_*F*_ is reached.

Enthalpy of combustion density given by equation (24) is defined in each discretization subinterval as 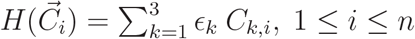, where 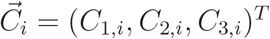. For Nash optimization, there are n objective functions *F*_Nash,*i*_, 1 ≤ *i* ≤ *n*, one for each subinterval. The input of *F*_Nash,*i*_ is the value of flux vector *β* on the *i*-th subinterval while its values on all other subintervals are fixed and denoted by 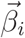, and the output of *F*_Nash,*i*_ is the enthalpy of combustion density on the ith subinterval, 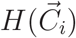. Thus,

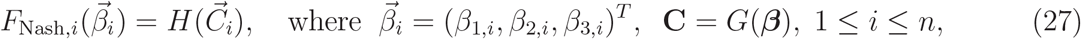

In the case of system optimization, there is one objective function *F*_Sys_ for the entire domain. The input for *F*_Sys_ is the entire flux vector *β*, and the output of *F*_Sys_ is the mid-point quadrature approximation of the *L*_1_ norm of the enthalpy of combustion density *H*(**x**). Thus,

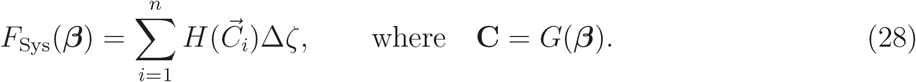

The discrete version of the influx constraints given by (20) and (23) on the *i*-th subinterval is

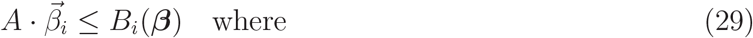

where

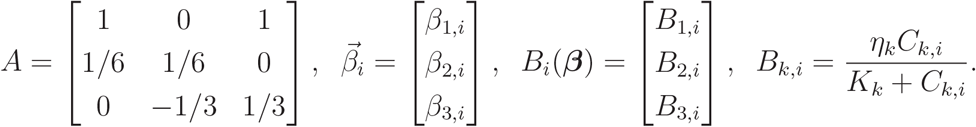

Since the right hand sides of the constraints (29) also depend on the unknowns *β*, a iterative procedure is applied to compute *β*. The following algorithm is used to find the optimal flux *β* and the associated chemical concentrations C for a fixed biofilm thickness.

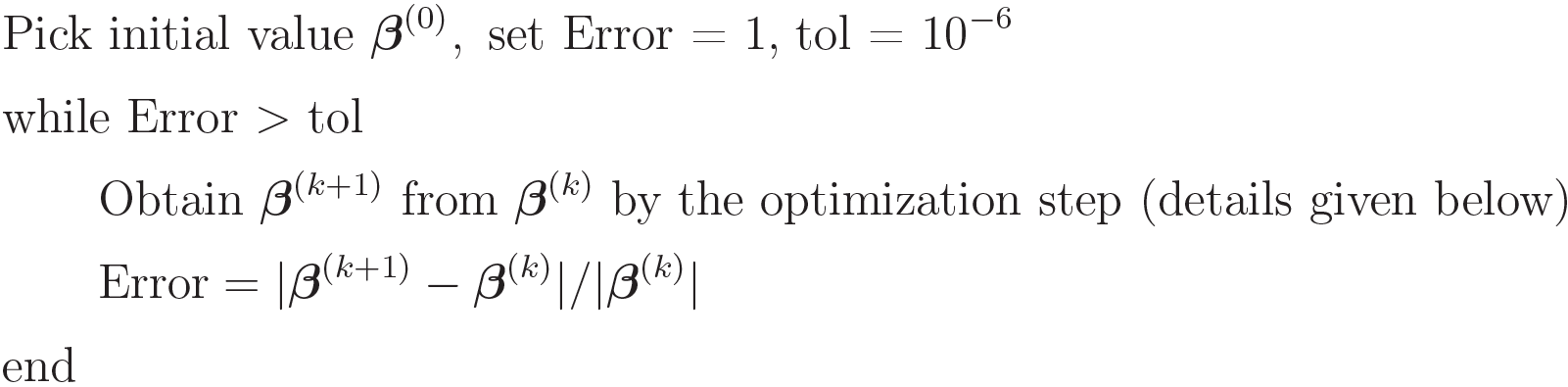

In the case of Nash optimization, the optimization step involves looping over all subintervals. Namely,

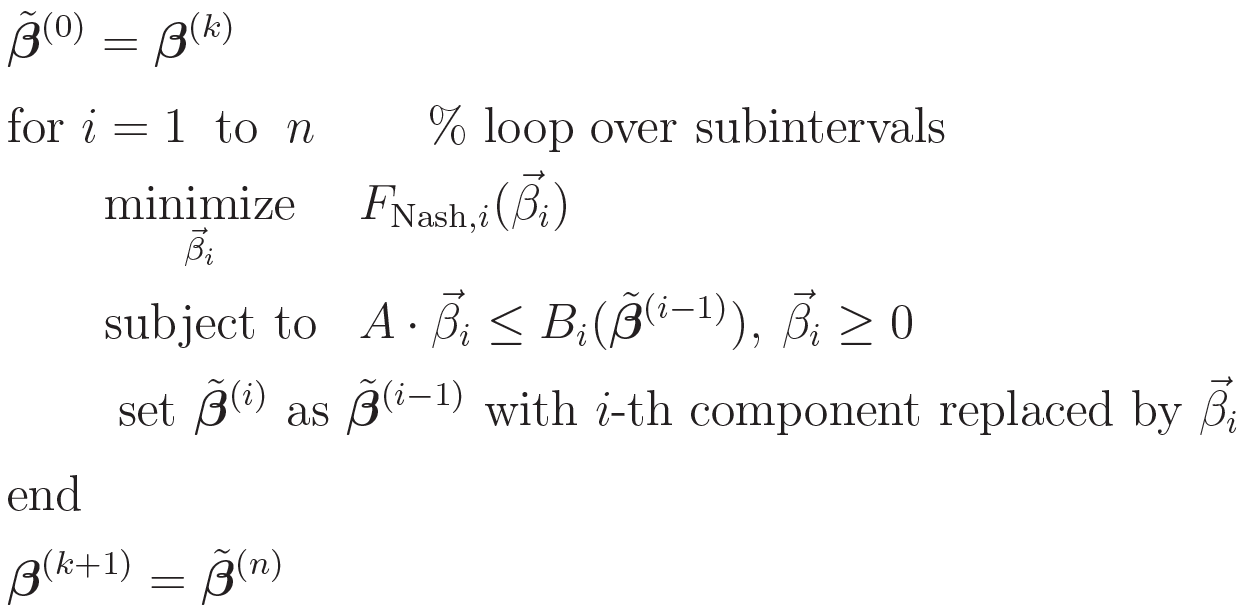

For System optimization, the optimization step is

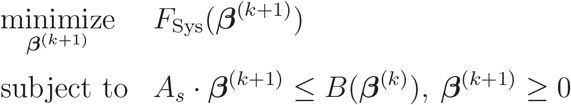

with

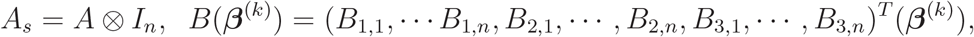

where ⊗ denotes the Kronecker product and *I*_*n*_ is the identity matrix of size *n*. In both cases, the constrained minimization problem is solved using the MATLAB function `fmincon`.

## 5 Results

### 5.1 Fixed Layers

We begin with fixed time snapshots of thin and thick biofilm layers, see Figure 3. In each, we consider a biofilm layer of fixed size with microbial metabolism as described in Fig. 1 and Section 3.1 at steady state. Oxygen, glucose, and lactate diffuse through the layer and exchange internally/externally as described in Section 3.2, and microbes at each spatial location locally optimally allocate their available resources, meaning that chemical flux into the local microbes is allocated to the three available metabolic pathways (glucose respiration, glucose fermentation, and lactate fermentation) in such a way as to minimize local exterior (to cells) enthalpy of combustion, i.e., computations reflect Nash equilibria based on environmental chemical concentrations as measured through enthalpy of combustion. Note: we have also used, in place of minimization of enthalpy of combustion, maximization of biomass production rate (18) as optimization objective, with little change in results. For systems in which growth rate and ebthalpy reduction are less closely linked, this might not be the case.

**Figure 3:**
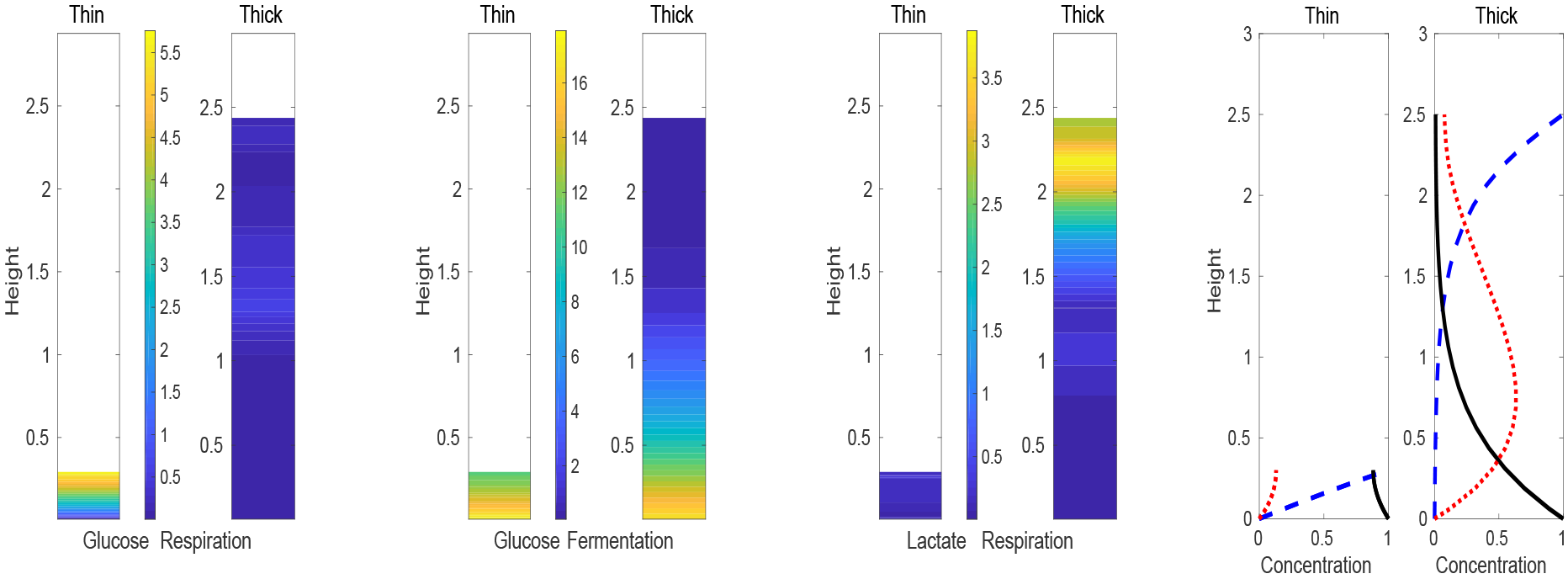
Snapshots of thin and thick biofilms. Bar plots show amplitudes of the elementary flux modes – each pair has the thin biofilm on the left and the thick biofilm on the right. Color bar in between indicates amplitude. Far left: glucose respiration. Middle left: glucose fermentation. Middle right: lactate respiration. Far right: line plots show chemical concentrations in, respectively, the thin and thick biofilm snapshots. (Blue dashed) oxygen concentration. (Black solid) glucose concentration. (Red dotted) lactate concentration. Note that the thin biofilm shows a profile combining glucose respiration and fermentation. The thick biofilm predominantly ferments glucose to lactate near the *z* = 0 interface and respires lactate near the *z* = 2.5 interface.

In the thin layer, relatively little consumption of resources occurs so that glucose penetrates fully. The same is essentially true for oxygen, though there is diffusive transport from the “air” boundary (*C*_1_|_*z*=*L*_ = 1) to the “tissue” boundary (*C*_1_|_*z*=0_ = 0) so that oxygen concentration is forced to 0 at *z* = 0 by the boundary condition. We see, notably, pathways for both respiration and fermentation of glucose are active throughout. Lactate respiration is mostly absent as oxygen is used preferentially for glucose respiration. Note that the model predicts glucose fermentation even in the presence of oxygen, consistent with some observations. This occurs here as a consequence of a mismatch between maximal uptake rates for oxygen and glucose; after allocation of all available oxygen influx to respiration, excess capacity for influx of glucose is directed to fermentation.

By contrast in the thick biofilm computation, we observe predominantly glucose fermentation metabolism in a lower sublayer where oxygen is unable to penetrate, and lactate respiration in an upper sublayer where oxygen is available but glucose is not. Note that lactate is produced in the lower layers and transferred to the upper layer by diffusive transport where it is consumed. Thus glucose respiration is mostly absent because glucose is largely depleted before it reaches the oxygen rich region. Similar profiles have been observed in single biofilm observations [41].

### 5.2 Growing Biofilm

Next, we consider a growing biofilm, see Figure 4, allowing the biofilm height to change according to (22). The left three panels of Fig. 4 show both the biofilm height as well as elementary flux mode amplitudes in time (horizontal axes are time, vertical cross-sections roughly correspond to the bar graphs in Fig. 3. Note that at early times, the distribution of fluxes in the biofilm is similar to that seen in the thin biofilm snapshot, Fig. 3, and at later times, the fluxes in the biofilm transitions to a distribution similar to that seen in the thick biofilm snapshot, Fig. 3.

**Figure 4:**
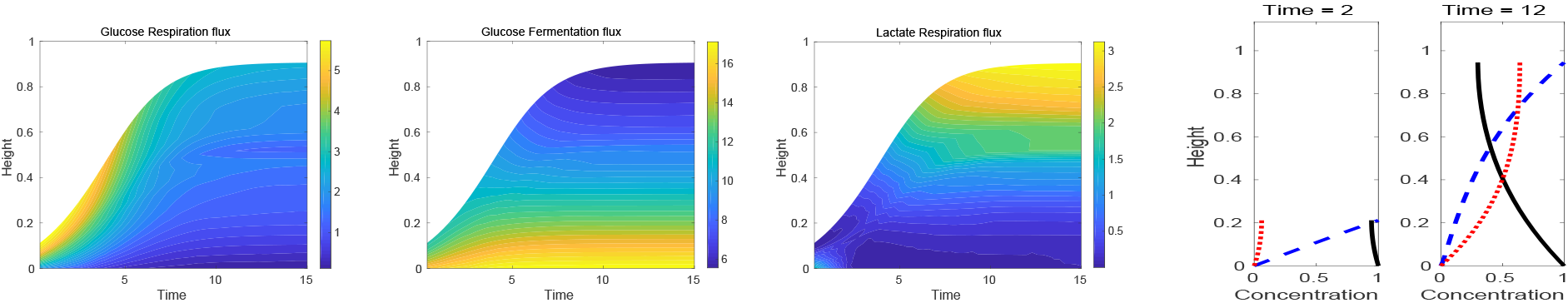
Growing biofilm. Left three panels are amplitudes of the three elementary flux modes, horizontal axes are time, vertical axes are height; height 0 is the substratum (the “tissue” interface), and the boundary between shaded and unshaded areas is top of the biofilm (the “air” interface). Shading indicates amplitude of flux, unshaded regions are exterior to the biofilm. Far left: flux through the glucose respiration pathway. Middle left: flux through the glucose fermentation pathway. Middle right: flux through the lactate fermentation pathway. Far right: plots show external chemical concentrations at *t* = 2 and *t* = 12: (blue dashed) oxygen concentration, (black solid) glucose concentration, (red dotted) lactate concentration. Note, at early times, the flux profile resembles that of a thin biofilm as in Fig. 3 dominated by a combination of glucose respiration and glucose fermentation. At later times, the flux profile resembles that of a thick biofilm as in Fig. 3 dominated by glucose fermentation near the “tissue” interface where oxygen levels are low and dominated by lactate respiration near the “air” interface where oxygen levels are high. At intermediate times, where the flux profile changes from thin biofilm to thick biofilm, the flux of new biomaterial also changes, reflecting the system-wide drop in metabolic efficiency and slowing of biofilm growth.

Observe that growth rate slows at the transition between thin and thick regimes, where glucose respiration is replaced by glucose fermentation to lactate combined with lactate respiration. In our simplified model, the latter is not significantly less efficient than the former. However, there is penalty paid for diffusive transport of lactate from where it is produced (near *z* = 0) to where it is consumed (near *z* = *L*). This transport penalty increases as biofilm height *L* increases, eventually approaching a cost at which growth and loss balance.

### 5.3 Nash vs System Optimization

To illustrate the importance of choice of optimization measure, we present an example involving a “mutant” organism which has lost the ability to respire lactate, see Figure 5. That is, the mutant can only respire glucose or ferment it to lactate. In this reduced metabolism, glucose fermentation is unfavorable in the sense that energy in lactate, the fermentation product, becomes lost to the system. Thus, if the system were to somehow coordinate its metabolism over the entire biofilm, for example via signalling of some sort, it might choose to avoid fermentation.

**Figure 5:**
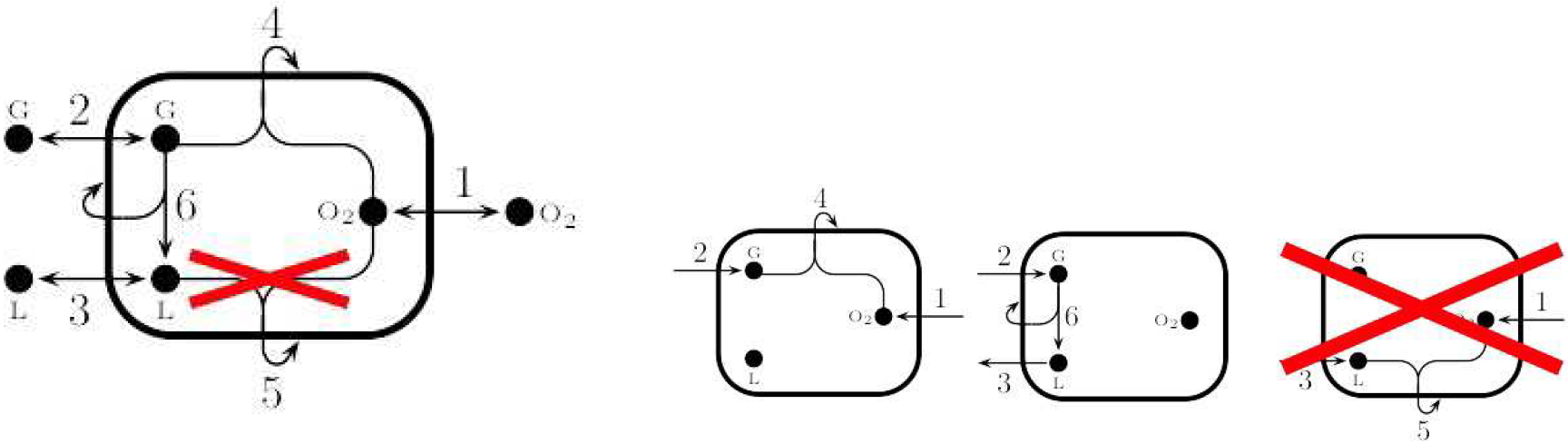
Reduced metabolism, compare to Fig. 1, without ability to respire lactate. Stoichiometric matrix is the same in (15) except with column 5 deleted. (Left) Metabolic model with suppressed reaction crossed out. (Right) Elementary flux modes for the metabolic model on the right, identical to those shown in Fig. 1 except that the lactate respiration pathway is unavailable.

We consider again a fixed time biofilm snapshot and compute the metabolic distribution to minimize enthalpy of combustion, as always, but using two different measures of optimality: (1) Nash optimality (as applied in other computations in this subsection) where metabolism is chosen based on local, pointwise, enthalpy without regards to non-local consequences, and (2) System optimality (used only here) where metabolism is chosen based on total system enthalpy (L_1_ norm). In the second case, microbes in different locations cooperate to produce a globally efficient biofilm.

See Figure 6 for an example computation. Note that the metabolic distributions are qualitatively different for the two optimization measures. In the Nash case, glucose is immediately consumed as it enters the biofilm from the substratum *z* = 0 via respiration above where oxygen is available and fermentation below where oxygen is depleted. In the System case, fermentation is highly unfavorable in comparison to respiration, so glucose is allowed to be diffusively transported toward the top of the biofilm where, despite the transport penalty, more oxygen results in increased respiration and overall decrease in system wide enthalpy. Looking at a comparison of the local enthalpy density in Fig. 6, we see that the Nash system chooses a small local advantage from fermenting glucose to lactate near the substratum leading, though, to (comparatively) significant excess enthalpy density above from unused lactate.

**Figure 6:**
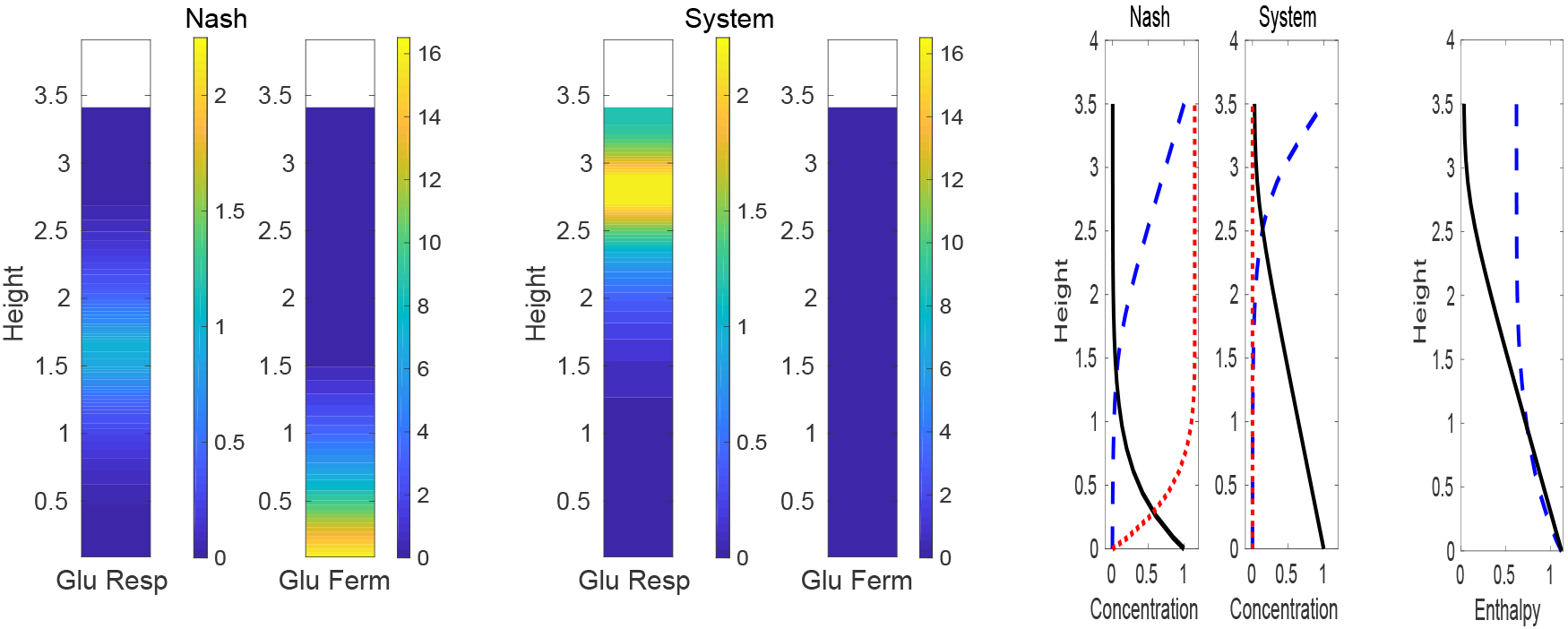
Snapshots of Nash-optimized and System-optimized biofilms with lactate respiration inhibited as in Fig. 5. Bar plots show amplitudes of the active elementary flux modes. Left: Nash optimized glucose respiration and fermentation flux amplitude. Middle: System optimized glucose respiration and fermentation flux amplitude. Right: first two line plots show chemical concentrations for, respectively, the Nash and System optimizations, with (blue dashed) oxygen concentration, (black solid) glucose concentration, (red dotted) lactate concentration. Third line plot shows density of enthalpy of combustion in UNITS of (blue dashed) Nash optimization and (black solid) System optimization. Note that the Nash optimized biofilm is dominated by glucose fermentation near the *z* = 0 interface with a small region above of where remaining glucose is respired, while, in the System optimized biofilm, glucose is transported diffusively to a respiration zone near the top of the biofilm. The plot of enthalpy density shows that there is a small local advantage to fermenting glucose near *z* = 0 at the cost of a large increase in global enthalpy due to unused lactate.

We remark that a contrast between Nash and System equilibrium is clear in this mutant system lacking lactate respiration capability. In the full metabolism, though, including lactate respiration capability, Fig. 1, the two equilibria are similar somewhat coincidentally: Nash optimized results are already reported in Fig. 3, the System optimization results (not shown) are similar. In particular, it is not perhaps obvious why System optimization chooses to ferment glucose near the bottom of the biofilm. The reason to do so is to minimize effect of the diffusive transport penalty. System optimization chooses to ferment glucose near the substratum and then diffusively transport lactate to the upper, oxygenated region, paying the diffusion transport penalty in lower enthalpy lactate rather than in glucose.

### 5.4 Data Integration

In some situations, additional data is available beyond metabolic capabilities, and in such cases it may be desirable to incorporate this information into a model. For example, given a set of observations of a microbial system and a hypothesized set of metabolic capabilities for its microbes, how do or don’t the observations constrain the actual metabolic activity, i.e., constraint what the microbes are doing and where they are doing it? This is an inverse problem mathematically, and difficult generally, but useful. A natural additional question: can we identify more or less helpful additional measurements, that is, measurements that have more or less constraining impact on our model results? This latter constitutes something of a sensitivity analysis.

Short of exploring these questions in depth here, we propose a method to begin moving in those directions by modifying the optimization objective (24) to account for additional known data. Particularly, if concentration data is known or inferred at a certain given locations *y*_*ℓ*_, 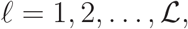, then we look to minimize using an augmented energy function of the form

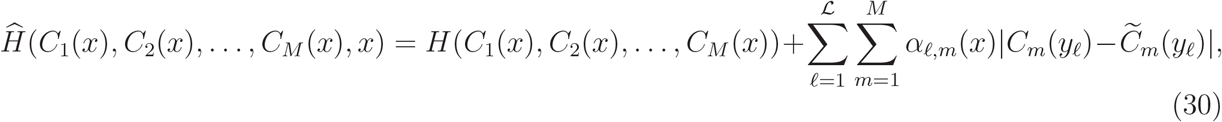

where 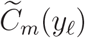 are the additional concentration data to be integrated and *α*_*ℓ,m*_(*x*) are weights (set to zero where data is not available). For example, see below, we may have additional information on concentrations at the boundary of a biofilm and wish to understand how that might constrain the metabolic profile inside the biofilm. By varying the magnitude of the weights *α*_*ℓ,m*_, the relative importance of the two components of 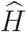 can be varied, though we do not explore this possibility here. Note that time dependence can also be allowed in (30) but is not included here for simplicity.

To illustrate, we employ a synthetic example based on the full metabolism (1) (with lactate respiration capability, designated LR positive) and the reduced metabolism (5) (without lactate respiration capability, designated LR negative), as follows: we first compute biofilm snapshot concentration profiles for the LR negative metabolism (5), identical in fact to the Nash optimized solutions shown in Fig. 6, and pretend that these profiles are actually an actual “real-life” experimental system from which we are able to extract only certain restricted bits of data via measurements of some sorts. Next, we provide that limited data, in this case some bits of information on boundary concentrations at the top *z* = 2.5 for oxygen and/or lactate, to the fully capable LR positive model (1). That is, we pretend to take a measurement of oxygen and lactate concentrations at the top of a “real-life” biofilm using the computed LR negative profiles as values for those measurements, then feed them to the LR positive model. The aim is to investigate how additional “data” taken from the “real-life” LR-negative profiles and provided to the LR-positive solver through objective (30) might impact predictions of the full, LR-positive model.

For all computations we use the same set of boundary conditions including, in particular, conditions (26) at the top of the biofilm layer. We provide “measured data” from the LR negative profiles to the LR positive model as follows. For lactate we can suppose that in addition to the no-flux condition on lactate at the top boundary, we also can measure the lactate concentration there (taken to be the LR negative concentration at *z* = 2.5). For oxygen we suppose that in addition to given the Dirichlet condition at the top, we also can measure the oxygen flux there (taken to be the top boundary oxygen flux from the LR negative computation). Mathematically, using optimization of (30), we try to force an additional lactate Dirichlet condition to the already imposed Neumann one, and/or for oxygen we try to force an additional Neumann condition to the already imposed Dirichlet one.

We consider the following cases for fitting:

1. LR positive without any additional LR negative data (LRP),
2. LR positive supplemented by the upper interface oxygen flux value “measured” from the LR negative profile (LRP+Oxy),
3. LR positive supplemented by the upper interface lactate concentration value “measured” from the LR negative profile (LRP+Lac),
4. LR positive supplemented by both the upper interface lactate concentration and oxygen flux values “measured” from the LR negative profile (LRP+Oxy+Lac).

As described, in case 1 we use boundary conditions as already imposed in (25)-(26) while in cases 2-4 we supplement those boundary conditions with the extra boundary data as follows. Prescription of glucose concentration is implemented numerically by setting the glucose concentration at the upper boundary node. Prescription of oxygen flux is equivalent to prescribing the derivative of oxygen concentration. Numerically this amounts to setting the oxygen concentration at the first discretization node into the domain from the boundary (oxygen concentration at the boundary node is already fixed by the Dirichlet boundary condition). In terms of a discretized version of (30), these amount to supplying 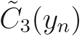 and/or 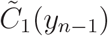 where *y*_*n*_ is the location of the top discretized point in the biofilm. Augmented objective (30) becomes

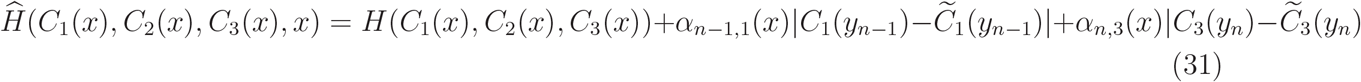

For weights we use *α*_*n*−1,1_ = *α*_*n*,3_ = 0.7 or 0 depending on the case.

See Figure 7 for concentration profiles for LR negative (left) and LR positive (right) profiles for reference. Note the most evident difference is in the lactate concentration profiles. For both LRN and LRP, lactate is produced in approximately the same region near the lower biofilm boundary. However, the LR-positive biofilm respires it above in the region where oxygen is available, while the LR-negative biofilm is unable to respire it at all. The aim below is to supplement the LR-positive computations with the extra data described above in order to, using an optimization objective (31), investigate how the new predictions resemble those for the LR-negative system particularly with respect to lactate concentration profile.

**Figure 7:**
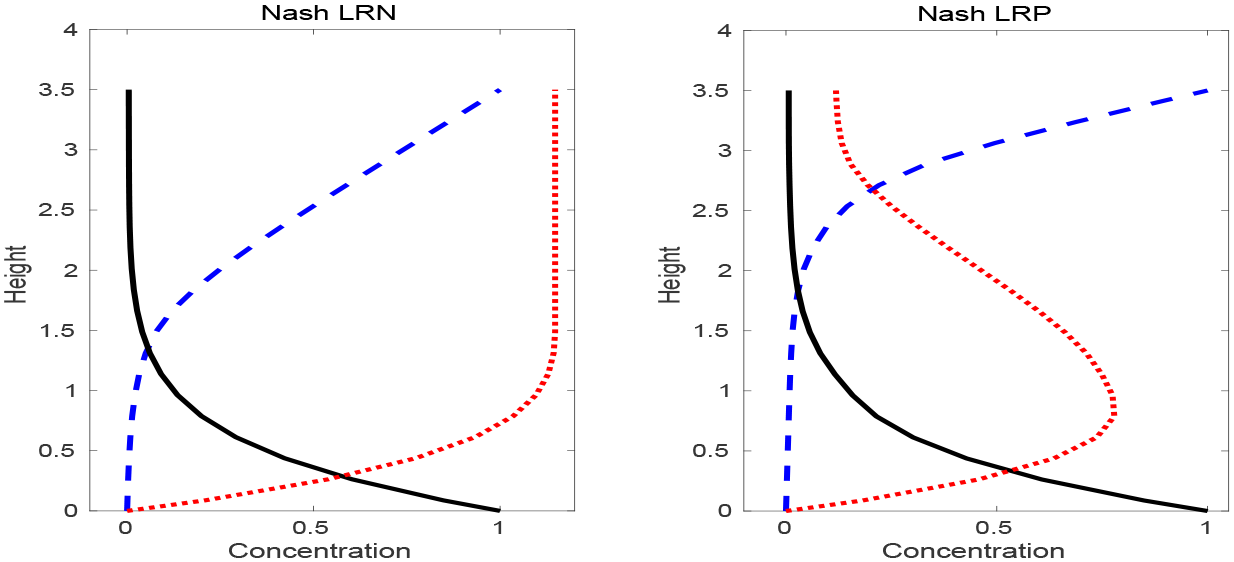
Substrate concentrations in the lactate respiration negative biofilm (left). and the lactate respiration positive biofilm (right). Nash optimization is applied for both. (Blue dashed) oxygen concentration. (Black solid) glucose concentration. (Red dotted) lactate concentration. LRN = lactate respiration-negative. LRP = lactate respiration-positive.

We show predicted concentration profiles under fitting cases 1-4 in Figure 8 columns 1 4, respectively, First, for reference again, column 1 compares LR-positive (without additional data) profiles to LR-negative profiles. The effects of adding oxygen boundary-flux data to (31) (*α*_*n*−1,1_ = 0.7, *α*_*n*,3_ = 0) are shown in column 2. Observe that the extra information about oxygen flux has little effect on predicted profiles. Though one might to hypothesize that net oxygen use (here largely determined by oxygen influx at the top boundary) is important and hence the extra oxygen flux data would be informative, in this example there is in fact relatively little difference in total oxygen use between the LR negative and positive systems so that the extra data has little effect on predictions. In contrast see column 3 where the LR positive system is augmented with lactate concentration data from the top boundary (*α*_*n*−1,1_ = 0, *α*_*n*,3_ = 0.7). Perhaps surprisingly, this extra information results in very close fits. Note the lactate concentration fits in the bottom row of Fig. 8. Both LR negative and LR positive systems generate lactate from glucose near the bottom of the biofilm where oxygen concetration is low (lower left subplot of Fig. 8). But it seems that forcing the LR positive system to, at the top boundary of the biofilm, match lactate concentration with the LR negative system effectively disallows lactate loss via respiration. Finally, for completeness, column 4 has plots from case 4, adding both the oxygen and lactate boundary data (*α*_n−1,1_ = *α*_*n*,3_ = 0.7), showing little change from the column 3 profiles.

**Figure 8:**
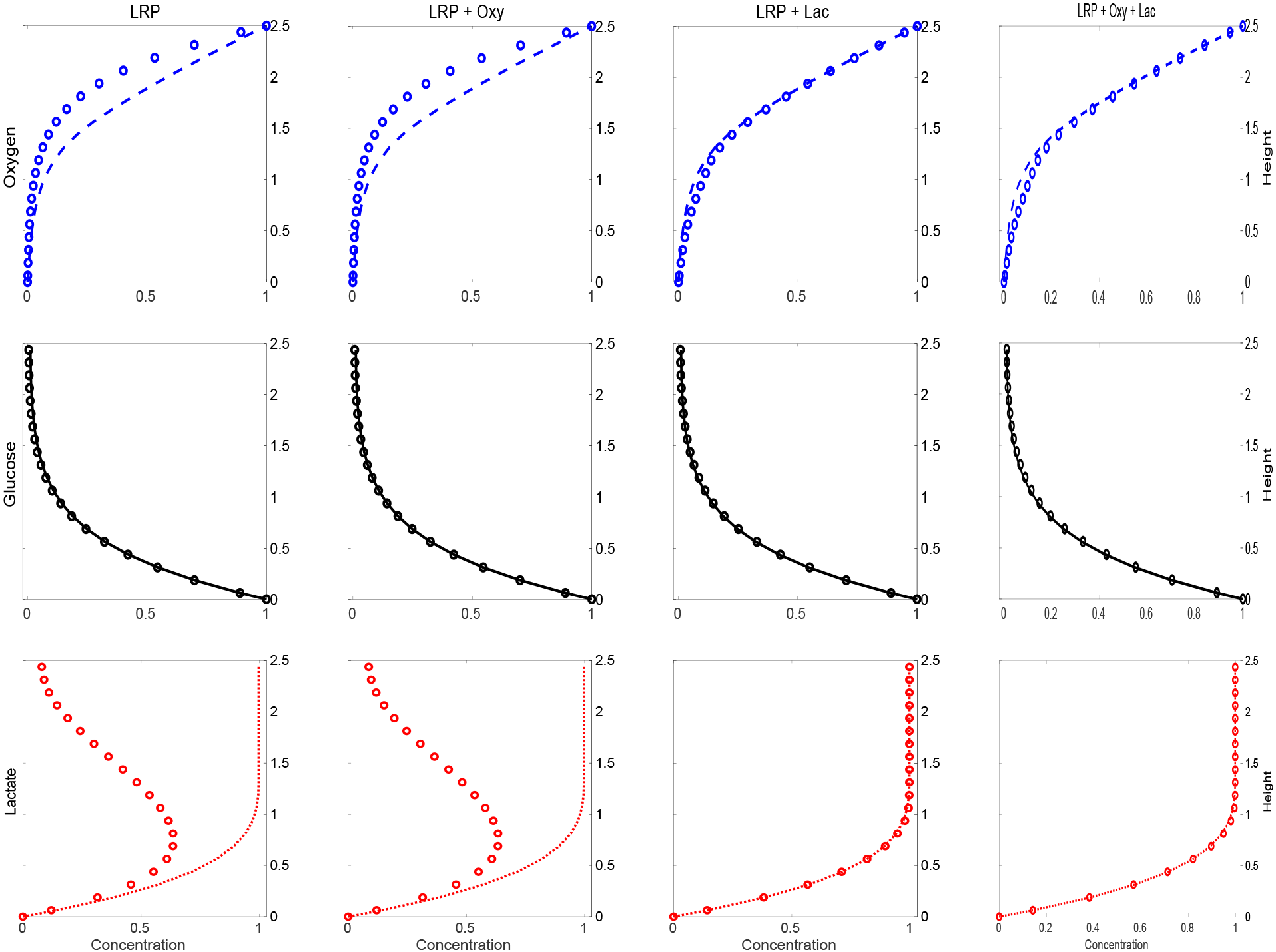
Comparisons between LR negative concentration profiles and augmented LR positive concentration profiles. Top row: oxygen. Middle row: glucose. Bottom row: lactate. First column: LR negative vs LR positive. Second column: LR negative vs LR positive + extra oxygen flux data. Third column: LR negative vs LR positive + extra glucose concentration data. Fourth column: LR negative vs LR positive + extra oxygen flux data + extra glucose concentration data. In each plot, the line style curve is the LR negative concentration profile and the circle symbols are the augmented LR positive concentration profile. Note that oxygen data augmentation does not improve the augmented LR positive predictions, while glucose data (with or without the oxygen data) does.

The quality of concentration fits does not extend to pathway flux amplitude fitting, see for example Figure 9 where we show profiles of lactate respiration flux amplitudes in the various cases 1-4 of the augmented LR positive system. The noise in the predicted pathway flux amplitudes is a consequence of the underdetermined linear relation between pathway fluxes *β*_*j*_ and exchange fluxes *e*_*k*_, see equations (20) which have a one dimensional null space; recall that exchange fluxes determine chemical concentrations through equations (21). That is, there are many different combinations of pathway fluxes which can lead to the same exchange fluxes. Thus fitting concentration data is indeterminant. To illustrate, the middle panel of Fig. 9 shows different realizations of the lactate respiration flux predicted by the LR positive system augmented by lactate boundary data (glucose respiration and fermentation pathway fluxes not shown); each realization however results in concentration profiles nearly identical to those seen in Fig. 9, right panel.

**Figure 9:**
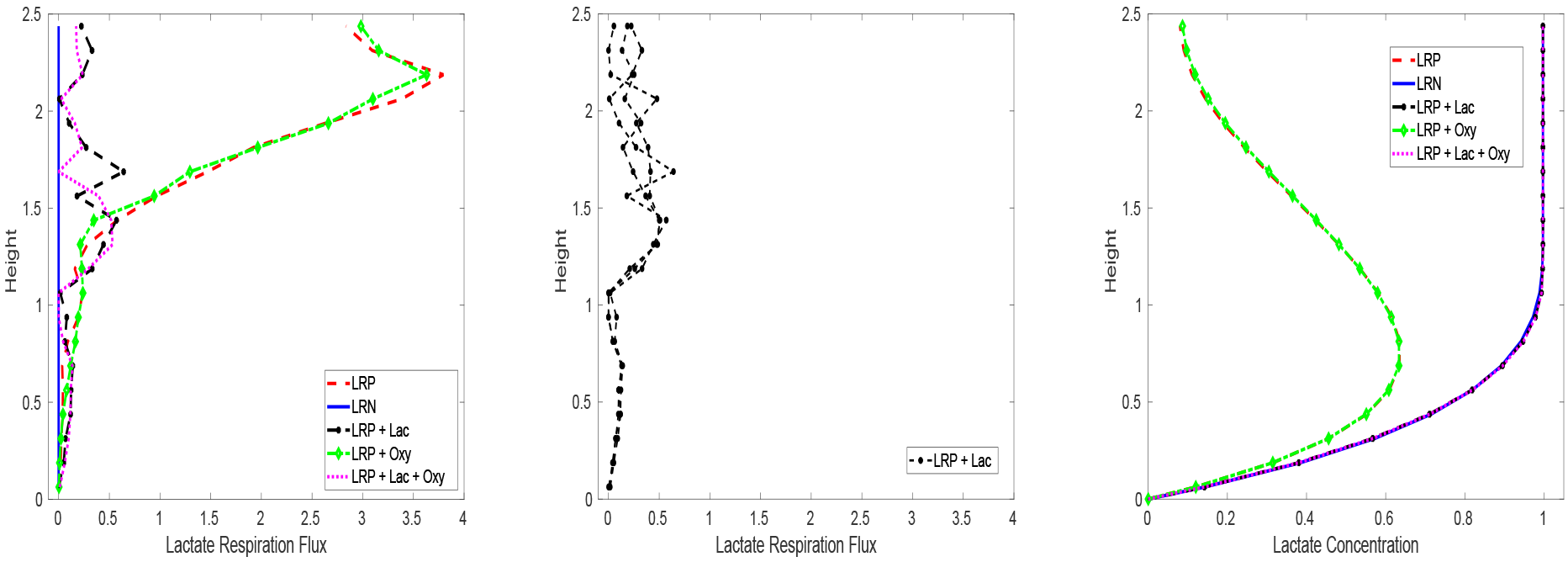
Amplitude of lactate respiration pathway flux for fitting cases LRP, LRP+Oxy, LPR+Lac, LPR+Oxy+Lac, see left panel. The solid blue line at 0 is the amplitude of lactate respiration pathway flux for the LR negative system. Middle panel: amplitude of lactate respiration pathway flux for three different realizations of the same LRP+Lac system. The noise is a consequence of indeterminancy in the solution. Noise also appears in all other pathway flux profiles. Right panel: for reference, lactate concentration profiles resulting from pathway fluxes (left and middle panels) are shown. The curves for LRP and LRP+Oxy lie nearly on top of each other, as do the curves for LRP+Lac, LRP+Lac+Oxy, and LRN. All three of the realizations shown in the middle panel result in the nearly the same LRP+Lac fit in the right panel.

The significance of the close match in columns 3 and 4 of Fig. 8 ahould not be overstated as both the PR negative and PR positive profiles arise from the same computational system except with different metabolic models. That is, the “data” is generated from a similar system as that used for fitting. Nevertheless, this example demonstrates the ability for easily connecting to experimental data and design. The pathway amplitude noise seen in Fig. 9 is also illustrative, as it is likely generally the case the concentration data is not sufficient uniquely determine the pathway flux profile.

## 6 Discussion

Population level models of microbial communities are burdened by often difficult to measure kinetics and regulatory parameters. The numbers of such parameters can rapidly increase as demands, fueled by availability of increasingly detailed molecular data, are made for more detailed descriptions of microbial community function. The result can be uncertainty in a high dimensional parameter space. However, at the cellular level and based on a steady state assumption, the metabolic modeling community has already independently recognized and developed methodologies to mitigate kinetic parameter overload and regulatory ambiguity. Our aim here has been to propose a procedure to marry steady state metabolic methods to population scale modeling, and to do so in a general fashion that allows flexibility in the mix and matching of different strategies both at the cellular and population levels. At the same time, as we couple cellular-scale metabolic models to the large, community and environmental scale, we have been able to introduce large scale environmental information back to the cellular level. Though not addressed here, it should further be possible to incorporate molecular information beyond genomics-derived metabolic maps.

Steady state metabolic modeling itself generally requires some selection principle for choosing an identified solution from an underdetermined set of possible solutions. We can employ standard strategies like maximization of biomaterial production. However, one potentially useful feature of the multiscale approach we propose is the possibility of naturally coupling larger scale environmental conditions to this selection, including thermodynamic or empirical data, and at local or non-local scale. Along the same lines, integration and inversion of data is also natural, either from large scale to small scale or vice-versa.

We have illustrated our methodology in the context of a one dimensional biofilm model, a moderately complicated system mathematically, but expect that the principles extend to more (e.g. multidimensional, multispecies biofilms in fluid or viscoelastic fluid environments) or less (e.g. batch or chemostat reactors) complex setups, or indeed to microbial communities beyond biofilms specifically. To change, essentially, replace the left-hand sides of (10) and (12) with different system-scale physics. Indeed, as long as the system time scale is long compared to the time scale for metabolisms to approach steady state, then the multiscale approach presented here seems applicable. In the absence of a separation of time scales, validity of steady state metabolic modeling might be questioned.

On the metabolic side, we have used a kinetics model characterized by elementary flux modes. Any other flux-driven model platform could be used instead. Further, we have supposed a heavily simplified, single organism metabolism. We have no reason to expect that moderately more complicated metabolisms cannot be readily substituted. However full (e.g. genome scale) metabolic models combined with more complex systems, like biofilms, would likely pose computational challenges requiring more attention to efficient numerical algorithms.

## Acknowledgements

The authors wish to acknowledge support from NSF/DMS awards 1517100 and 1516951 as well as The Fields Institute.

## References

[1] T. Bjarnsholt, M. Alhede, M. Alhede, S.R. Eickhardt-Sørensen, C. Moser, M. Kühl, Ø.P. Jensen, and N. Høiby, The in vivo biofilm. Trends in Microbiology 21, 466–474 (2013).

[2] D.E. Moormeier, J.L. Endres, E.E. Mann, M.R Sadykov, A.R. Horswill, K.C. Rice, P.D. Fey, and K.W. Bayles, Use of microfluidic technology to analyze gene expression during Staphylococcus aureus biofilm formation reveals distinct physiological niches. Appl. Environ. Microbiol. 79, 3413–3424 (2013).

[3] J. Dockery, I. Klapper, Finger formation in biofilm layers, SIAM J. Appl. Math. 62, 853–869 (2002).

[4] J.E. Bailey, Complex biology with no parameters, Nature Biotech 19, 503 (2001).

[5] A. Bordbar, J.M. Monk, Z.A. King, B.Ø. Palsson, Constraint-based models predict metabolic and associated cellular functions, Nat. Rev. Genet. 15, 107âĂŞ120 (2014).

[6] D. Calvetti, J. Heino, E. Somersalo, K. Tunyan, Bayesian stationary state flux balance analysis for a skeletal muscle metabolic model, Inverse Prob Imag 1, 247–263 (2007).

[7] D. Calvetti, E. Somersalo, Quantitative in silico analysis of neurotransmitter pathways under steady state conditions, Front Endocrinol 4, article 137 (2013).

[8] R.P. Carlson, Decomposition of complex microbial behaviors into resource-based stress responses, Bioinformatics 25, 90–97 (2009).

[9] W.G. Characklis, Fouling biofilm development: A process analysis, Biotech. Bioeng. 23, 1923âĂŞ1960 (1981).

[10] J Chen, J.A. Gomez, K. Höffner, P. Phalak, P.I. Barton, M.A. Henson, Spatiotemporal modeling of microbial metabolism, BMC Syst Biol 10:21 (2016).

[11] B.L. Clarke, Stoichiometric network analysis, Cell Biophys. 12, 237–53 (1988).

[12] N.G. Cogan, Two-fluid model of biofilm disinfection, Bull Math Biol 70, 800–819 (2008).

[13] N.G. Cogan, J.P. Keener, The role of the biofilm matrix in structural development, Math. Med. Biol. 2, 147–166 (2004).

[14] J.A. Cole, L. Kohler, J. Hedhli, Z. Luthey-Schulten, Spatially-resolved metabolic cooperativity within dense bacterial colonies, BMC Syst Biol 9, DOI 10.1186/s12918-015-0155-1 (2015).

[15] B. D’Acunto, L. Frunzo, I. Klapper, M.R. Mattei, Modelling multispecies biofilms including new bacterial species invasion, Math Biosci 259, 20–26 (2015).

[16] B. D’Acunto, L. Frunzo, I. Klapper, M.R. Mattei, P. Stoodley, Mathematical modeling of dispersal phenomenon in biofilms, to appear, Math Biosci, (2018).

[17] N.C. Duarte, M.J. Herrgård, B.Ø. Palsson, Reconstruction and validation of Saccharomyces cerevisiae iND750, a fully compartmentalized genome-scale metabolic model, Genome Res 14, 1298–1309 (2004).

[18] R Duddu, D.L. Chopp, B. Moran, A twoâĂŘdimensional continuum model of biofilm growth incorporating fluid flow and shear stress based detachment, Biotech Bioeng 103, 92–104 (2009).

[19] H.J. Eberl, R. Sudarsan, Exposure of biofilms to slow flow fields: The convective contribution to growth and disinfection, J Theor Biol 253, 788–807 (2008).

[20] J.S. Edwards, R.U. Ibarra, B.Ø. Palsson, In silico predictions of Escherichia coli metabolic capabilities are consistent with experimental data, Nat Biotechnol 19, 125–130 (2001).

[21] W.R. Harcombe, W.J. Riehl, I. Dukovski, B.R. Granger, A. Betts, A.H. Lang, G. Bonilla. A. Kar, N. Leiby, P. Mehta, C.J. Marx, D. Segrè, Metabolic resource allocation in individual microbes determines ecosystem interactions and spatial dynamics, Cell Reports 7, 1104–1115 (2014).

[22] M.A. Henson, P. Phalak, Byproduct cross feeding and community stability in an in silico biofilm model of the gut microbiome, Processes 5, doi:10.3390/pr5010013 (2017).

[23] K.A. Hunt, J.P. Folsom, R.L. Taffs, R.P. Carlson, Complete enumeration of elementary flux modes through scalable demand-based subnetwork definition, Bioinformatics 30, 1569–1578 (2014).

[24] N. Jayasinghe, A. Franks, K.P. Nevin, R. Mahadevan, Metabolic modeling of spatial heterogeneity of biofilms in microbial fuel cells reveals substrate limitations in electrical current generation, Biotech. J. 9, 1350–1361 (2014).

[25] I. Klapper, J. Dockery, Role of Cohesion in Material Description of Biofilms, Phys. Rev. E 74, 031902 (2006).

[26] I. Klapper, J. Dockery, Mathematical description of microbial biofilms, SIAM Rev. 52, 221–265 (2010).

[27] I. Klapper, B. Szomolay, An exclusion principle and the importance of mobility for a class of biofilm models, Bull Math Biol 73, 2213–2230 (2011).

[28] N. Klitgord, D. Segrè, Environments that induce synthetic microbial ecosystems, PLoS Comput. Biol. 6, Article e1001002 (2010).

[29] A. Khodayari, C.D. Maranas, A genome-scale Escherichia coli kinetic metabolic model k-ecoli457 satisfying flux data for multiple mutant strains, Nat Commun 7, 13806 (2016).

[30] J.-U. Kreft, Biofilms promote altruism, Microbiology 150, 2751–2760 (2004).

[31] P.J. Linstrom and W.G. Mallard, Eds., NIST Chemistry WebBook, NIST Standard Reference Database Number 69, National Institute of Standards and Technology, Gaithersburg MD, 20899, doi:10.18434/T4D303, (retrieved May 25, 2018).

[32] J. Liu, A. Prindle, J. Humphries, M. Gabalda-Sagarra, M. Asally, D.D. Lee, S. Ly, J. Garcia-Ojalvo, G.M. Süel, Metabolic co-dependence gives rise to collective oscillations within biofilms, Nature 523, 550–554 (2015).

[33] R. Mahadevan, J.S. Edwards, F.J. Doyle III, Dynamic flux balance analysis of diauxic growth in Escherichia coli, Biophys. J. 83, 1331–1340 (2002).

[34] Algorithmic Game Theory, N. Nisan, T. Roughgarden, É. Tardos, V.V. Vazirani, Eds., Cambridge University Press, Cambridge (2007).

[35] J.D. Orth, I. Thiele, B.Ø. Palsson, What is flux balance analysis?, Nat Biotechnol 28, 245–248 (2010).

[36] P Phalak, J Chen, R.P. Carlson, M.A. Henson, Spatiotemporal metabolic modeling of a chronic wound biofilm consortium, IFAC-PapersOnLine 49–26, 032âĂŞ037 (2016).

[37] C. Picioreanu, M.C.M. van Loosdrecht, J.J. Heijnen. Effect of diffusive and convective substrate transport on biofilm structure formation: a two-dimensional modeling study. Biotech. Bioeng. 69, 504–515 (2000).

[38] B. Polizzi, O. Bernard, M. Ribot, A time-space model for the growth of microalgae biofilms for biofuel production, J. Theor. Biol. 432, 55–79 (2017).

[39] N.D. Price, J.L. Reed, B.Ø. Palsson, Genome-scale models of microbial cells: evaluating the consequences of constraints, Nat Rev Microbiol 2, 886–897 (2004).

[40] H. Qian, D.A. Beard, Thermodynamics of stoichiometric biochemical networks in living systems far from equilibrium, Biophys Chem 114, 213–220 (2004).

[41] S.A. Rani, B. Pitts, H. Beyenal, R.A. Veluchamy, Z. Lewandowski, W.M. Davison, K. Buckingham-Meyer, P.S. Stewart, Spatial patterns of DNA replication, protein synthesis, and oxygen concentration within bacterial biofilms reveal diverse physiological states, J Bacterial 189, 4223–4233 (2007).

[42] T.D. Scheibe, R. Mahadevan, Y. Fang, S. Garg, P.E. Long, D.R. Lovley, Coupling a genome-scale metabolic model with a reactive transport model to describe in situ uranium bioremediation, Micrab Biatechnal 2, 274–286 (2009).

[43] J. Schellenberger, B.O. Palsson, Use of randomized sampling for analysis of metabolic networks, J Bial Chem 284, 5457–5461 (2009).

[44] C.H. Schilling, S. Schuster, B.Ø. Palsson, R. Heinrich, Metabolic pathway analysis: basic concepts and scientific applications in the post genomic era, Biotech Pragr 15, 296–303 (1999).

[45] R. Schuetz, L. Kuepfer, U. Sauer, Systematic evaluation of objective functions for predicting intracellular fluxes in Escherichia cali, Mal Syst Bial 3, 119 (2007).

[46] S. Schuster, S. Hilgetag, On elementary flux modes in biochemical reaction systems at steady state, J Bial Syst. 2, 165âĂŞ182 (1994).

[47] H.-S. Song, D.G. Thomas, J.C. Stegen, M. Li, C. Liu, X. Song, X. Chen, J.K. Fredrickson, J.M. Zachara, T.D. Scheibe, Regulation-structured dynamic metabolic model provides a potential mechanism for delayed enzyme response in denitrification process, Frant Micrabial 8, 01866 (2017).

[48] P.S. Stewart, Diffusion in biofilms, J. Bacteriol. 185 1485âĂŞ1491 (2003).

[49] P.S. Stewart, M.J. Franklin, Physiological heterogeneity in biofilms, Nat. Rev. Micrabiol. 6, 199210 (2008).

[50] R. Taffs, J.E. Aston, K. Brileya, Z. Jay, C.G. Klatt, S. McGlynn, N. Mallette, S. Montross, R. Gerlach, W.P. Inskeep, D.M. Ward, R.P. Carlson, In silico approaches to study mass and energy flows in microbial consortia: a syntrophic case study, BMC Sys Biol 3, 114 (2009).

[51] G.D. Tartakovsky, A.M. Tartakovsky, T.D. Scheibe, Y. Fang, R. Mahadevan, D.R. Lovley, Pore-scale simulation of microbial growth using a genome-scale metabolic model: Implications for Darcy-scale reactive transport, Adv Water Resour 59, 256–270 (2013).

[52] P. Unrean, F. Srienc, Metabolic networks evolve towards states of maximum entropy production, Metab Eng 13, 666–673 (2011).

[53] J.B. van Klinken, K.W. van Dijk, FluxModeCalculator: an efficient tool for large-scale flux mode computation, Bioinformatics 32, 1265–1266.

[54] S.J. Wiback, R. Mahadevan, B.Ø. Palsson, Reconstructing metabolic flux vectors from extreme pathways: defining the *α*-spectrum, J. Theor. Biol. 224, 313–324 (2003).

[55] T. Zhang, B. Pabst, I. Klapper, P. S. Stewart, General theory for integrated analysis of growth, gene, and protein expression in biofilms, PloS one 8, (2013).

[56] A.R. Zomorrodi, C.D. Maranas, OptCom: a multi-level optimization framework for the metabolic modeling and analysis of microbial communities, PLOS Comp Biol 8, e1002363 (2012).

